# Duplications and retrogenes are numerous and widespread in modern canine genomic assemblies

**DOI:** 10.1101/2023.10.31.564742

**Authors:** Anthony K. Nguyen, Matthew S. Blacksmith, Jeffrey M. Kidd

**Affiliations:** Department of Human Genetics, University of Michigan, Ann Arbor, Michigan; Department of Computational Medicine and Bioinformatics, University of Michigan, Ann Arbor, Michigan

**Keywords:** read-depth, alignment, segmental duplication, retrocopy, genome

## Abstract

Recent years have seen a dramatic increase in the number of canine genome assemblies available. Duplications are an important source of evolutionary novelty and are also prone to misassembly. We explored the duplication content of nine canine genome assemblies using both genome self-alignment and read-depth approaches. We find that 8.58% of the genome is duplicated in the canFam4 assembly, derived from the German Shepherd Dog Mischka, including 90.15% of unplaced contigs. Highlighting the continued difficulty in properly assembling duplications, less than half of read-depth and assembly alignment duplications overlap, but the mCanLor1.2 Greenland wolf assembly shows greater concordance. Further study shows the presence of multiple segments that have alignments to four or more duplicate copies. These high-recurrence duplications correspond to gene retrocopies. We identified 3,892 candidate retrocopies from 1,316 parental genes in the canFam4 assembly and find that approximately 8.82% of duplicated base pairs involve a retrocopy, confirming this mechanism as a major driver of gene duplication in canines. Similar patterns are found across eight other recent canine genome assemblies, with multiple metrics supporting the high-quality of the mCanLor1.2 wolf assembly constructed using PacBio HiFi reads. Comparison between the wolf and other canine assemblies found that approximately 92% of retrocopy insertions are shared between assemblies. By calculating the number of generations since genome divergence, we estimate that new retrocopy insertions appear, on average, in 1 out of 3,514 births. Together, our analyses illustrate the impact of retrogene formation on canine genomes and highlight the variable representation of duplicated sequences among recently completed canine assemblies.

**Significance:** Duplications are highly influential on evolution, but are commonly misassembled, especially in lagging genomic groups like canines. We assessed nine canine assemblies for duplication presence, and found enrichment for acrocentric regions, misattribution of duplications to unplaced contigs, and the presence of short, high-recurrence duplications. Investigating further, we find high numbers of retrocopies retaining hallmarks present in the canine assemblies, and determine a rate of novel retrocopy insertion at 1 in 3,514 births.

## Introduction

Gene duplication is a major driver of evolution, but correctly resolving duplications in genome assemblies remains challenging. Gene duplication is a form of structural variation, which refers to chromosomal alterations that are typically larger than 1000 bases (1 Kb) in size, and includes other changes such as insertions, deletions, inversions, and other copy number changes [1]. Copy number gains that affect genes are a key mechanism in the evolution of novel gene functions and the emergence of gene families. Duplicated genes are associated with a number of core biological processes, including development, transcriptional regulation, and environmental response [2].

Gene duplications arise through two means: the reverse transcription and integration of RNA transcripts, or the duplication of DNA segments. Duplications on the DNA-level can arise via whole genome duplications, in which the entire genome is replicated. Whole genome duplications have occurred multiple times in evolutionary history and have contributed to increased genome complexity in vertebrates [3]. Alternatively, DNA-level duplications may be smaller and affect only a segment of the genome. These segmental duplications may encompass genes, giving rise to additional copies of genes and the expansion of related gene families.

Segmental duplications are typically defined as non-repetitive sequences longer than 1 Kb that are found in more than one copy in a haploid genome [4]. This definition differentiates duplications from common repeats, such as LINEs (Long INterspersed Element) and SINEs (Short INterspersed Element), which proliferate throughout genomes through distinct mechanisms.

Segmental duplications are further divided into three types based on the relative locations of duplicates: tandem duplications, in which duplications appear adjacent to one another; intra-chromosomal dispersed duplications, which is when duplications occur on the same chromosome but with an intervening sequence; and inter-chromosomal dispersed, when duplication copies occur on separate chromosomes. The intervening sequence between related duplicates can include genes and is susceptible to additional duplication or deletion [5]. DNA duplications likely first arise as a result of replication-based errors including break-induced replication and template switching mechanisms [6]. Once formed, additional copies can be gained or lost due to nonallelic homologous recombination [7]. No matter their type, when these copies differ in count between individuals or populations they are referred to as copy number variants (CNVs).

Gene duplications may also be caused by a retrotransposition-associated mechanism, in which mRNAs are reverse transcribed and inserted into the genome at new locations [8]. The resulting duplications, known as retrocopies, lack introns and show the hallmarks of retrotransposition, including a poly(A) tail and flanking target site duplications [9]. While retrocopies refer to any derivation of a gene transcript from retrotransposition, whereas the term retrogene implies functionality, both terms will be used interchangeably within this study [10]. The retrocopies may be full-length or truncated at their 5’ end. Over time, mutations in retrocopies may accumulate that disrupt gene function creating pseudogenes [11]. However, retrogenes may be functional; several have been documented to have associations with important biological processes, including immune function, metabolism, environmental response, physical appearance, and disease, in both humans and canines [10, 12]. Gene conversion and changes in pseudogene sequence can also give rise to actual and novel function, altering the name from pseudogenes into functional retrogenes. Retrocopies commonly occur in dogs [13, 14] and confound the interpretation of genetic variation [15].

Novel developments in sequencing technology have allowed for the rapid and thorough analysis of thousands of samples. Despite this, systematically identifying and analyzing duplications remains challenging. To investigate CNVs in samples, the most common method of analysis is to look for an increase in read-depth, typically obtained from whole genome sequencing, relative to a reference genome assembly. Duplications can also be found by aligning a genome reference to itself to create a genome self-alignment that identifies sequences present multiple times [16]. When constructing such genome self-alignments, it is critical to first filter out common repeats since the proliferation of repeats overwhelms the alignment process.

There are several challenges associated with properly assembling duplicated regions. Assembly errors may lead to collapsed duplications in which duplicated sequences are only represented once in an assembly [17]. This commonly occurs at the end of assembled contigs due to a failure to correctly assemble sequences spanning the duplication. Such assembly errors can also give rise to chimeric contigs that have been incorrectly assembled as a result of mis-joining non-adjacent segments [18]. False duplications can also arise due to misassembly. This can occur when contigs created from divergent alleles are mistakenly represented as being duplicated [19–21]. Correctly assembling duplications is further complicated by the diploid nature of most analyzed genomes. As a result of challenges in correctly assembling duplicated sequences, the duplication content of a genome assembly may not be an accurate representation of the true duplication content of the analyzed genome. Assemblies derived from short sequencing reads are plagued by incorrect representation of duplicates [20]. Longer sequencing reads coupled with new methodologies for haplotype-resolved assembly [22, 23] promise to alleviate these concerns resulting in a more accurate representation of genome content and structure.

The canine remains a powerful system for genetics and genomics. The domestic dog is in the unique position of having co-evolved alongside humans, while simultaneously experiencing human-driven extreme artificial selection for over ten thousand years [24]. The first canine reference genome was released in 2005, derived from Tasha the Boxer [25]. This was further refined into canFam3.1, released in 2014, also based on Tasha [26]. The canine reference genome was assessed for copy number variation across multiple studies. Initial studies estimated that approximately 4.21% of the canine reference genome is composed of recent segmental duplication, including 841 genes in predicted segmental duplications [27]. Array-CGH data showed that many duplications were copy-number polymorphic [28]. Additional studies of canine CNVs have linked these variations to conditions such as ACL rupture [29] and breed-specific morphological structure [30]. Other studies have concluded that CNVs likely did not have major effects on canid domestication, with suggestions that dogs and wolves had similar proportions of CNV loci [31, 32].

Since the release of canFam3.1, multiple assemblies of dogs and wolves have been generated using long-read technology [33–40]. These new assemblies have helped elucidate novel genes, regulatory elements, and variants, while revealing new aspects of canine genome biology. These assemblies have been created using a variety of sequencing technologies (Oxford Nanopore and PacBio) and algorithms. However, limited analyses of the duplication content of these assemblies have been performed, with most description focused on specific loci, such as the amylase region, or other regions of suspected misassembly [41]. To address the lack of analysis of duplications in these long-read genomes, we aim to characterize the duplication content of different recently published long-read canine genomes and assess patterns of gene duplication found throughout canines. We apply two computational analyses to nine recently-released canine genome assemblies: genome assembly self-alignment, which involves the mapping of an assembly to itself in order to find matching duplicated segments [16] and read-depth analysis, which searches for regions of unusually high coverage in Illumina sequencing data [31, 41]. We further perform a search for retrocopy insertions in each genome assembly. We find notable and large regions of duplicated sequence that is not properly represented in existing dog assemblies, describe polymorphic duplications that differ between canine assemblies, and identify retrocopies as a major driver of gene duplications in canine genomes.

## Results

### Duplications identified by self-alignment of the Mischka genome include segments with multiple duplicated copies

The UU_Cfam_GSD_1.0 assembly (accession GCA_011100685.1), derived from a female German Shepherd Dog named Mischka, has extensive annotation, and serves as the reference genome used by the Dog10K consortium [14, 33, 42]. This genome assembly was constructed using PacBio Sequel sequencing and consists of sequence assigned to 39 assembled chromosomes (chr1-38 and chrX), the mitochondria (chrM), and 2,158 unplaced contigs not localized to a chromosome. Due to the importance of this reference to ongoing research, we first performed a detailed analysis of duplications in this assembly. Using genome assembly self-alignment, we identified 491,661 pairwise duplicated segments of 1,000 bp or greater with at least 90% sequence identity in the Mischka assembly. Of these, 122,521 segments are located in the 38 canine autosomes or chromosome X and 369,135 duplications are located on unplaced contig sequences. Of the duplicated segments on the assembled chromosomes, 50,629 have a paralog on an unplaced contig (41.03%). The largest single duplicated segment is nearly 1.8 Mb in length, located at chr8:74446738-76245570.

Many of the duplications detected on the assembled chromosomes are small; 46.62% (57,120/122,521) of duplicated segments are shorter than 2 Kb. Examination of the size distribution of duplicated segments smaller than 2 Kb revealed three notable peaks at approximately 1,250, 1,375, and 1,775 bp. Inspection of the duplication landscape on UCSC Genome Browser identified numerous high-recurrence duplications that contained many short alignments linking a locus with multiple paralogous segments **(Supplementary Figure 1)** [43].

We defined a high-recurrence segment as any area that contained at least four or more paralogous duplications smaller than 2.5 Kb, and identified 1,492 high-recurrence duplications, totaling approximately 3.0234 Mb. Overall, this makes up 3.095% of duplicated base pairs located by self-alignment. The segment with the largest number of copies is found at chr1:100216700-100219200, with 159 intersecting duplications. These high-recurrence duplications appear to be distributed throughout the Mischka genome.

Because the same locus can be included in multiple duplications at varying levels of sequence similarity, we merged segments together, resulting in 6,040 duplicated intervals in the assembled chromosomes and 1,940 duplicated intervals in the unplaced contigs (**Supplementary Figure 2, Supplementary Table 1 and 5**). Duplications occupy 96.89 Mb, or about 4.12%, of the assembled chromosomes. In sharp contrast, about 90.15% (115.8 Mb) of the sequence on unplaced contigs is duplicated. In total, including unplaced contigs, 8.58% of the Mischka assembly is duplicated according to genome assembly self-alignment.

Duplications are not distributed uniformly along canine autosomes. Of the 5,525 duplicated segments located on the autosomes, 193 (3.49%) overlap the first megabase of the beginning of the chromosome, while 172 (3.11%) are located within the last megabase of the autosomes. Relative to 1000 random permutations, this is a 2.5-fold increase for the first megabase (mean: 1.7%; max: 2.2%) and a 1.75-fold increase for the last megabase (mean: 1.6%; max: 2.2%).

### Duplications identified in Mischka using read-depth are not uniformly distributed

To complement duplications identified via genome assembly self-alignments, we identified duplications based on read-depth. Illumina sequencing reads derived from Mischka were aligned to the Mischka assembly and processed using fastCN [31]. Repetitive sequences are masked in the assembly prior to alignment and the read-depth profiles are used to estimate absolute copy number in windows consisting of 1,000 unmasked positions. We define read-depth duplications as segments consisting of four or more consecutive windows with an estimated copy number exceeding 2.5 and with a minimum size of 10 Kb, resulting in 1,573 segmental duplications defined by read-depth **(Supplementary Table 2 and 6)**. We find that approximately 60% (30.6 Mb) of unplaced contigs that contain at least four windows are duplicated. In comparison, 56.21 Mb (1033 segments) is duplicated on the assembled chromosomes, making up 2.39% of the 38 autosomes and X chromosome.

The duplications identified by read-depth range in size from 10 Kb to 1.6 Mb and have a mean size of 48 Kb and a median of 21 Kb. We find that 57 (6.1%) duplications are present in the first megabase of the autosomes, while 36 (3.8%) are located in the last 1 Mb of each autosome. Relative to random permutations, we find a 3.19-fold increase in duplication content in the first megabase of autosomes (mean: 1.9%; max: 3.5%), and a 2.1-fold increase in the last megabase (mean: 1.8%; max: 3.2%). The median copy number of all the duplications was 3.62, and the mean was 4.85. The maximum copy number of a given duplication was 78.3, located at chr27:33386819-33407975.

Most chromosomes have between 1% and 5% of their sequence annotated as a duplication by read-depth; however, 8.85% of chromosome 26 is duplicated. Examination revealed that a 2.4 Mb region encompassing 6 segmental duplications, located at chr26:25,387,954-27,841,554, accounts for this high duplication percentage. This region has a median estimated copy number of 9.5.

### Read-depth analysis and self-alignment show low levels of agreement

We compared the position of duplications identified by genome self-alignment and read-depth to search for concordance. To account for detection differences between the methods, we only considered duplications that were located on assembled chromosomes, greater than 15 Kb in size, have at least 95% identity, and were not located in the first megabase of an autosome. A total of 1,122 duplications identified by genome assembly self-alignment and 918 duplications identified by read-depth met these criteria **(Supplementary Table 1, 2, 5, 6)**.

We explored how many duplications in each set were found to be unique. By intersecting the self-alignment set and the read-depth set, we find that 528/1,122 self-alignment duplications (47.1%) and 504/918 read-depth duplications (54.9%) share any overlap. The union of these two sets contains approximately 77.78 Mb in total, 26.6 (34.2%) of which is duplicated in both sets. Self-alignment uniquely detects 29.88 Mb (38.41%) and read-depth uniquely detects 21.02 Mb (27.02%) **(Figure 1B)**. Despite only 35% agreement between the two methods, both analyses show similar trends at a chromosomal level, including an increase in duplication count on chr26 **(Supplementary Figure 3).**

**Figure 1.**
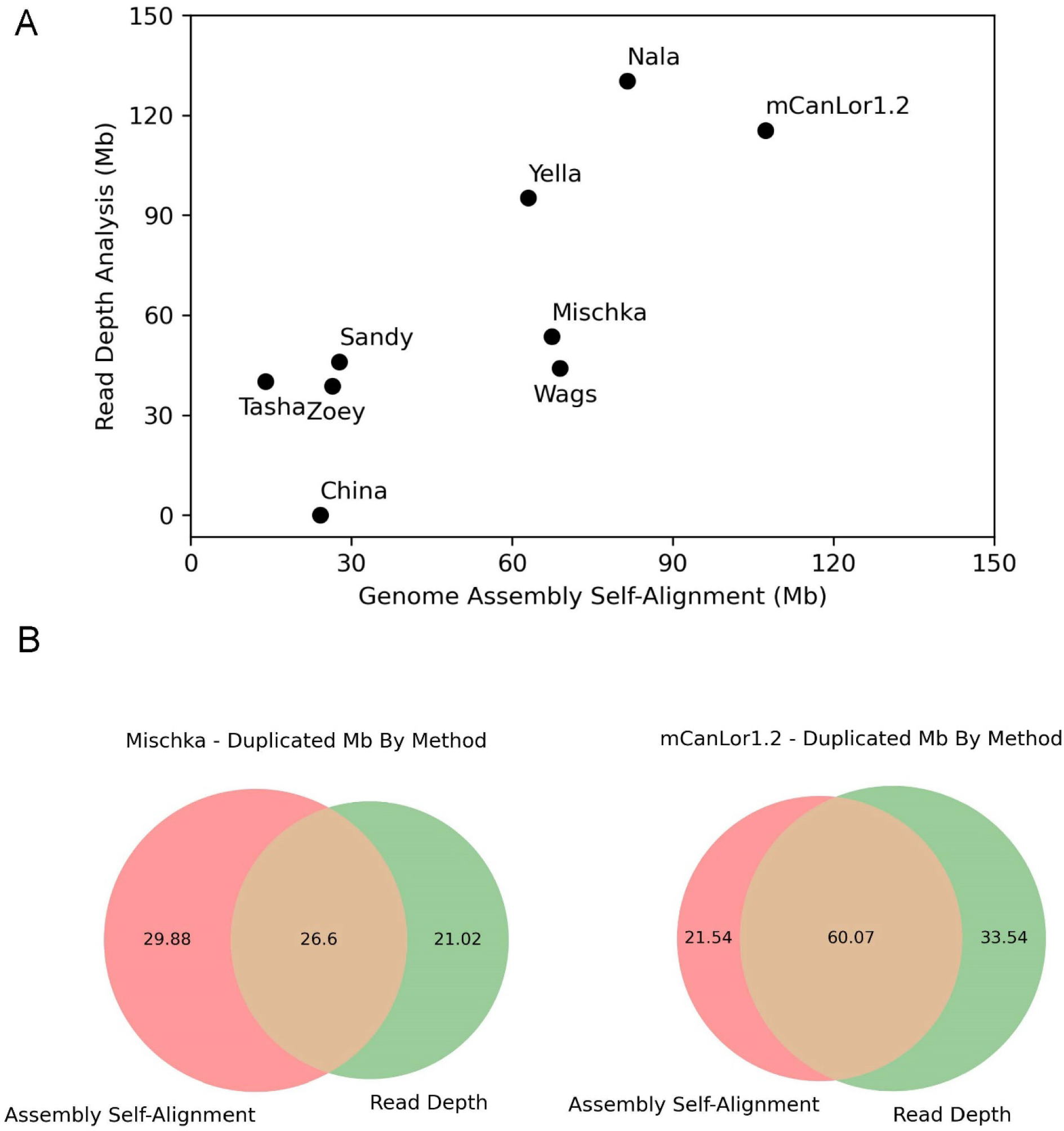
*Estimated duplication content of canine assemblies*. Duplication content was determined based on read-depth and genome assembly self-alignment. (A) The amount of each genome assembly detected as duplicated using each method is shown. Since the read-depth approach is best able to detect larger, more similar duplications, the regions detected by genome self-alignment were filtered to segments with 95% sequence identity at least 15 Kb in length. China lacks Illumina data and thus has no read-depth analysis. (B) A Venn Diagram showing the intersection of the two methods for the Mischka assembly and for the mCanLor1.2 assembly. Values are plotted in Mb.

### Duplication content varies across canine assemblies

We expanded our self-alignment and read-depth analysis to eight additional recently published canine assemblies **(Table 1**, **Figure 1A)** [33–40]. Most assemblies used some method of PacBio sequencing, predominantly Sequel, but the China and Yella assemblies used Oxford Nanopore (ONT) sequencing. We note that China and Wags are different Basenjis sequenced using different technologies by the same study. While the authors of that paper state that China is the better assembly, we exclude China from the read-depth analysis due to a lack of available Illumina data [40]. Notably, among the genomes is mCanLor1.2, a Grey wolf genome that serves as an outgroup to the other breed dog genomes [34].

**Table 1.** Canine Assemblies Investigated. Listed in each row is the name of the sample used for each assembly, the sample type, the N50 in megabases as reported by each publication, and the main sequencing approach used to construct the assembly as reported in each publication. Aside from Mischka/canFam4 at the top, the rest of the assemblies are ordered by their contig N50.

| Sample Name | Sample Type | N<sub>50</sub> (Mbp) | Primary Sequencing Technology | Accession Number |
| --- | --- | --- | --- | --- |
| Mischka (canFam4) | German Shepherd | 14.96 | PacBio Sequel | GCA_011100685.1 |
| Sandy | Dingo | 40.7 | PacBio Sequel/ONT PromethION | GCA_003254725.2 |
| China | Basenji | 37.76 | ONT PromethION | GCA_013276365.2 |
| mCanLor1.2 | Greenland Wolf | 34 | PacBio HiFi | GCA_905319855.2 |
| Tasha | Boxer | 27.45 | PacBio Sequel | GCA_000002285.4 |
| Nala | German Shepherd | 20.9 | PacBio Sequel | GCA_008641055.3 |
| Zoey | Great Dane | 4.72 | PacBio RSII | GCA_005444595.1 |
| Wags | Basenji | 3.13 | PacBio Sequel | GCA_004886185.2 |
| Yella | Labrador Retriever | 0.091 | ONT GridION | GCA_012045015.1 |

Canine assemblies have between 1.8% (Tasha) and 5.6% (mCanLor1.2) of their genome identified as duplicated by genome assembly self-alignment **(Table 1)**. Assemblies with a low duplication content (below 2.5%) include Sandy, Tasha, and Zoey, whereas assemblies with a high percentage of duplications (above 4%) include Mischka, mCanLor1.2, Nala, Wags, and Yella. A similar divide in duplication content among assemblies remains after filtering for duplications above 15 Kb and increasing the sequence identity minimum to 95%. By analyzing read-depth, we see a slight change, where mCanLor1.2, Nala, and Yella have the highest fraction of their genome annotated as duplicated. The number and duplication status of unplaced contigs differs widely among the assemblies **(Supplementary Table 1)**. The highest duplicated fraction is found in Mischka (90%) and Zoey (70%), with the others having less than 50% of their unplaced contigs duplicated.

Additionally, we explored 13,817 protein-coding genes for their duplication status as determined by read-depth. Illumina data from each sample was aligned to the Mischka assembly; China and Nala are removed from this analysis, due to lack of Illumina data and strong bias based on GC composition respectively **(Supplementary Figure 4)**. The copy number of each gene was then estimated using read-depth. We identify a total of 1,068 genes as duplicated in at least one canine; 222 of those genes are considered duplicated in all seven canines. Further analysis indicated that the most highly duplicated genes have no annotated function and are not associated with a named gene **(Supplementary Table 7).** We performed a Gene Ontology (GO) enrichment analysis of the 423 genes identified as duplicated based on read-depth from Mischka. Enriched functional categories include ‘intermediate filament organization’ (GO:0045109), ‘signal transduction’ (GO:0007165), ‘positive regulation of cellular process’ (GO:0048522), ‘multicellular organismal process’ (GO:0032501), and ‘negative regulation of cellular process’ (GO:0048523) **(Supplementary Table 9)** [44–46].

To explore the impact of PacBio HiFi reads on representing duplications, we compared duplications identified by assembly self-alignment and read-depth in the mCanLor1.2 assembly. Using the same filtering criteria as used in Mischka, we find that a total of 115.3 Mb is duplicated in the union of self-alignment and read-depth, a 1.5-fold increase over Mischka. We also find an increase in congruency between the two methods, as 60.07 Mb is duplicated in both the self-alignment set and the read-depth set, making up 52.11% of the total base pairs, a notable increase from the 34.2% overlap found in Mischka **(Figure 1C)**.

### Retrocopies proliferate throughout canine genome assemblies and show hallmarks of retrotransposition

While exploring self-alignment duplications, we found that high-recurrence duplications in the Mischka assembly often corresponded to the exons of transcripts found in other species, suggesting the presence of gene retrocopies **(Figure 2A)**. To confirm this, we identified candidate retrogenes in the Mischka assembly using BLAT. From an initial set of 19,972 multi-exon protein-coding genes, we identified 3,892 retrocopies from 1,316 parental genes, which includes 66 retrocopies that retained the full open reading frame of the parental gene **(Table 2, Supplementary Table 4)**. 3,658 (94%) of these retrocopies are located on assembled chromosomes. We found an enrichment of retrocopies on chrX, with a 1.18-fold increase over expectation from 1,000 random permutations **(Supplementary Figure 5).** Additionally, retrocopies that retained the full open reading frame of their parental gene had much greater sequence identity with the parent gene (median: 99.92%) vs. those that did not (median: 96.14%) (p-value: 3.81*10^-48^). (**Supplementary Figure 6).**

**Figure 2.**
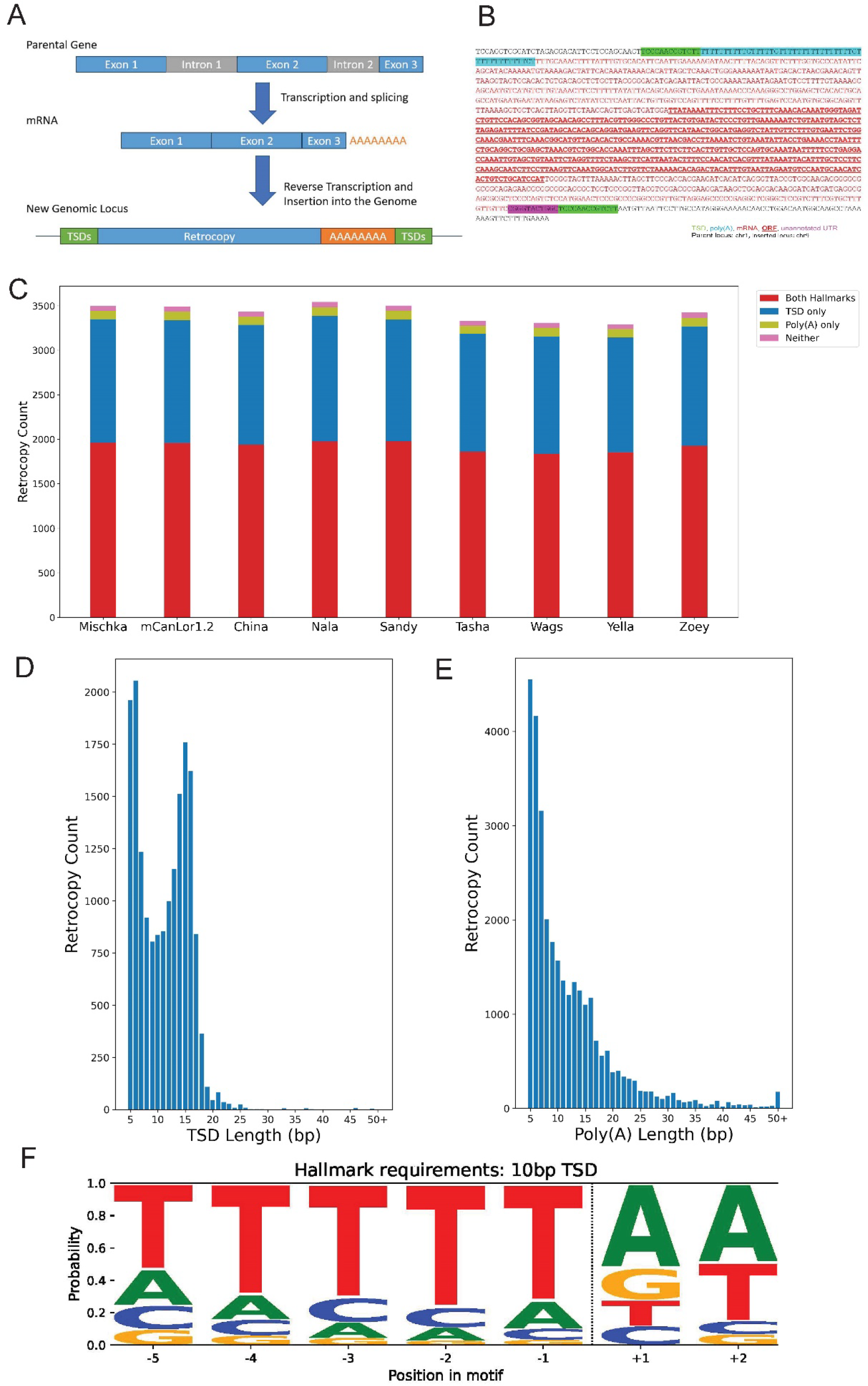
*Hallmarks of retrotransposition in detected retrocopies*. Analysis of retrocopy presence across canine assemblies is shown. A) A diagram depicting the key steps of retrocopy formation, including the presence of target site duplications and poly(A) tails at the insertion, is shown. B) The annotated sequence of a retrocopy from the TBPL1 parent gene, which is inserted in chr9 in the Mischka assembly, is shown. Highlighted sequence depicts the aforementioned hallmarks as indicated. This retrocopy retains the parental ORF, shown in bold. The short purple highlighted sequence corresponds to UTR sequence included in the retrocopy that is not part of the existing TBPL1 annotation. C) The distribution of how many retrocopies possess TSDs, poly(A) tails, both, or neither in each of the nine canine assemblies. D) A histogram of the TSD lengths found for retrocopies in the Mischka assembly. TSD lengths of 50 or greater have been grouped into one bin. E) A histogram of the lengths of poly(A) tails found for retrocopies in the Mischka assembly. Lengths of 50 or greater have been grouped into one bin. F) An examination of target site cleavage sequence for retrocopies in Mischka for retrocopies with at least a 10bp TSD (N=1615). It shows the sequence composition spanning 5 bases before and 2 bases after the inferred cut site.

**Figure 3.**
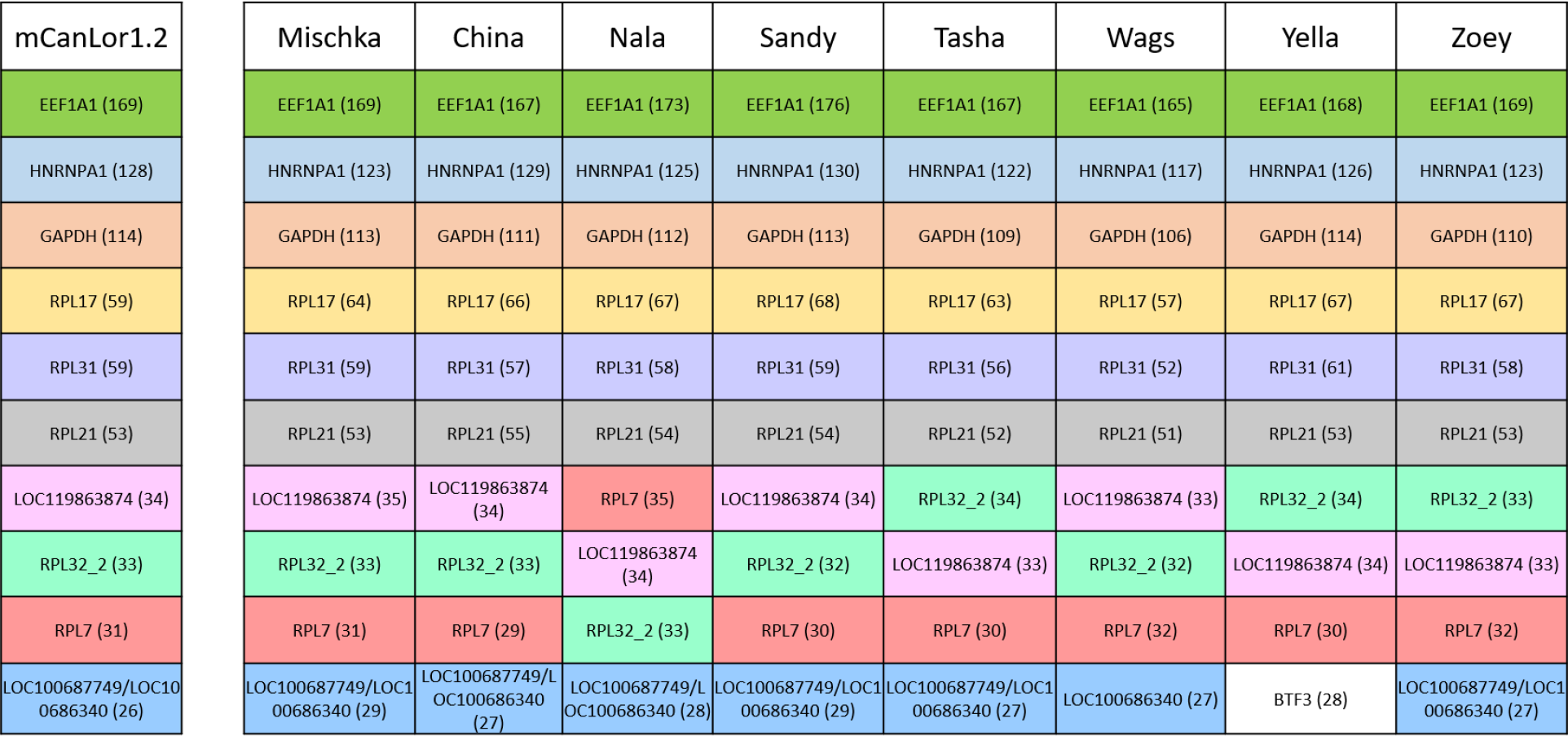
*Most frequent retrocopy parental genes*. Each column lists the 10 genes that give rise to the most identified retrocopies in each assembly. Retrocopies presented here are limited to the assembled chromosomes. The number of retrocopies from each parent are given in parentheses. The same genes are colored identically to one another to quickly determine presence of reoccurring genes. Only 11 parent genes are represented across all nine assemblies, with retrocopies from BTF3 found in the other assemblies at slightly decreased frequencies that removed it from the top ten.

**Table 2.**
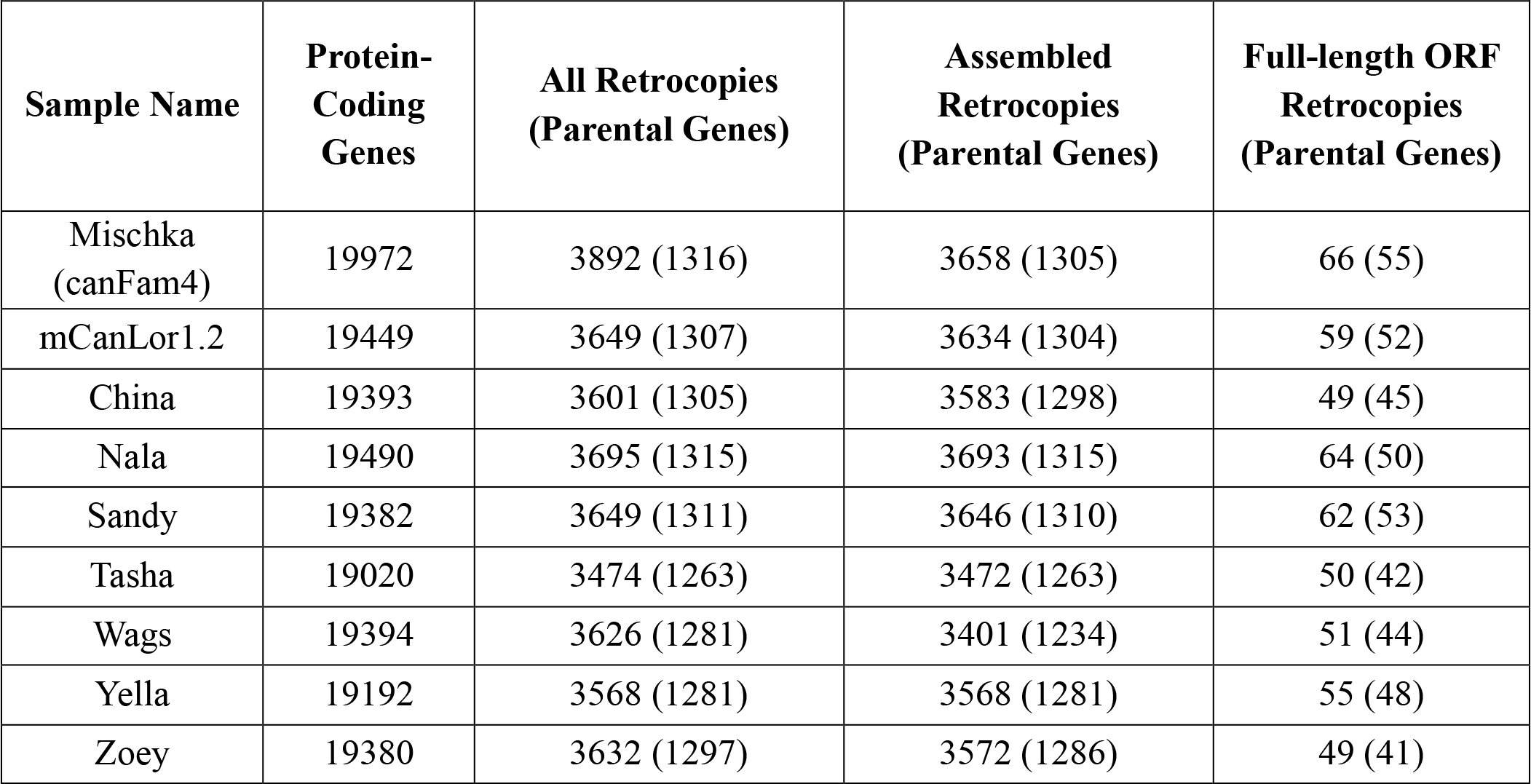
Retrocopies identified in each canine assembly. Column two gives the number of protein-coding genes considered as potential gene parents in each assembly. Retrocopies in column 3 include all chromosomes, including mitochondrial and unplaced contigs, and column 4 contains only the assembled chromosomes, chr1-38 + X. The total number of retrocopies and the number that retain the open reading frame of the parental gene are shown.

By intersecting high-recurrence duplications with annotated retrocopies, we determine that 1,080/1,492 (72.39%) high-recurrence duplications segments overlap with a retrocopy; in turn, 1,078/3,658 (29.47%) retrocopies on assembled chromosomes intersect with high-recurrence duplications. On a base-pair level, retrocopies located in highly duplicated segments make up a total of 4.35 Mb. Thus, multiple pair-wise alignments among retrocopies derived from the same parent gene account for the highly duplicated regions identified by genome self-alignment.

We additionally investigated how many retrocopies retained the signature hallmarks of retrotransposition, which are the presence of target site duplications (TSDs) and poly(A) tails [47] **(Figure 2B)**. Of the 3,496 retrocopies on assembled chromosomes in the Mischka genome that passed filtering, 3,442 (98.46%) show at least one of the two hallmarks **(Figure 2C)**. In total, 1,387 (39.67%) only have TSDs of at least 5 bp, 94 (2.69%) only have poly(A) tails, and 1,961 (56.09%) have both signatures. The mean length of poly(A) tails is 11.91 bases, and the mean length of TSDs is 11.19 bases, with 1082 having TSDs 10 bp or longer **(Figures 2D and 2E)**. All 66 retrocopies that retain the parental ORF have TSDs, while 20 also have a poly(A) tail.

Another known hallmark of retrotransposition is the presence of a consensus LINE-1 endonuclease recognition sequence, commonly read as TTTT/AA, where “/” marks the cleavage site. We searched for the presence of the expected consensus sequence across retrocopies and recovered the expected endonuclease recognition sequence **(Figure 2F)** [48–51]. We find that approximately 40% of retrocopies annotated with a TSD 5bp or longer contain the expected cut site sequence.

When analyzing only the retrocopy loci that are dimorphic among assemblies that passed filters (n=212), thus permitting a more precise definition of event boundaries based on comparison with the pre-integration empty site, we find a confirmed TSD of at least 5 bp or greater, with an average TSD length of 15.1 bp. Additionally, loci that possessed a confirmed TSD also possessed an endonuclease cleavage site **(Supplementary Figure 7).**

We expanded the retrocopy analysis to all other canine assemblies, using protein-coding genes converted from Mischka coordinates as queries. Retrocopy counts are broadly similar across assemblies (**Table 2, Supplementary Table 8**). Mischka’s slightly greater number of retrocopies appears to be accounted for by the presence of unplaced contigs, many of which correspond to duplicated sequence **(Supplementary Table 1).** Similar trends of identified TSDs and poly(A) tails are observed across the other assemblies, with over 98% of all identifies retrocopies showing at least one hallmark **(Supplementary Table 3)**. We also specifically analyzed retrocopies that retain the parental ORF for presence of hallmarks **(Supplementary Table 4)**. Over 96% of full-ORF retrocopies show at least one hallmark, usually presence of a target-site duplication.

The top ten retrocopy-producing parent genes were determined for each assembly **(Figure 4)**. There were only 11 parent genes present across the top ten lists of all assemblies, highlighting a high level of shared parental genes across the assemblies. Many of the top annotated genes are previously known retrocopy producers in human and/or canine (*GAPDH, HNRNPA1, EEF1A1, HMGN2* and the *RPL* genes) [52–56]. One gene, *BTF3*, was unique to the Labrador Retriever genome, Yella, but retrocopies from this gene are present at slightly decreased frequency in the other canine assemblies. Additionally, there were several instances where we could not determine a unique progenitor among related parental genes. Most notable among this group are LOC100686340 and LOC100687749, which are present in the top ten retrocopy producer lists of all canine assemblies except Yella and Wags, the latter of which only has LOC100686340 annotated.

To explore the function of common retrocopy progenitors, we performed a GO enrichment analysis of the most common parent genes in the Mischka assembly. There is one significant enriched GO category among the 50 parent genes in the Mischka assembly that had at least 10 retrocopies, ‘cytoplasmic translation’ (GO:0002181), which likely derives from the high number of ribosomal genes (RPL and RPS). There are 5 RPL genes in the top 10 parent genes, and 15 RPL or RPS genes in the top 50. We expanded this analysis to the 117 genes that had at least 5 retrocopies, which identified several enriched categories, including ‘ribosomal small subunit assembly’ (GO:0000028) and ‘cytoplasmic translation’ (GO:0002181) **(Supplementary Table 9)** [44–46].

Finally, we filtered retrocopies for any overlap with segmental duplications identified by read-depth. We find that the Mischka assembly has a high prevalence of retrocopies present in segmental duplications, with 466 compared to the range of 62-263 (avg: 126) found in other assemblies. We note that 162/466 (34.7%) are present on unplaced contigs, further highlighting Mischka’s duplication bias towards unplaced contigs. No other canine assembly has any retrocopies mapped to unplaced contigs, indicating that Mischka retrocopies are frequently present in segmental duplications that get assembled onto unplaced contigs, in line with the increased duplication rate present in the self-alignment data.

### Estimating the rate of retrocopy insertion

To identify the rate of retrocopy insertion, we compared five canine assemblies from diverse dogs (Mischka, Nala, China, Sandy, and Zoey) against mCanLor1.2. Only retrocopies located on the autosomal chromosomes that did not overlap a read-depth segmental duplication were considered. First, we assessed which retrocopies were shared across these six assemblies. After applying strict filtering, we found a total of 3,377 unique retrocopies across the six assemblies, 3,111 of which were located in all six canines (92.12%). 46 were unique to the mCanLor1.2 assembly, and the largest categories tended to be singletons **(Supplementary Figure 8, Supplementary Table 10)**. Mischka and Nala, both German Shepherds, have the fewest singletons, reflecting their similarity relative to the other assemblies.

We also searched for the presence of retrocopies in other Canines. We searched for 3,658 retrocopies from Mischka in the fox (VulVul2.2) and dhole (GWHAAAC00000000) genome assemblies [57, 58]. In total, 1923 retrocopies (52.2%) were found in the fox and/or the dhole genomes, 468 retrocopies (12.79%) were present in both the fox and the dhole, 549 were unique to fox (15%), and 906 were unique to the dhole (24.76%) **(Supplementary Table 11)**.

Consistent with their more recent origin, none of the 66 Mischka retrocopies with intact ORFs were found in either the fox or dhole assemblies. We also find that sequence similarity of retrocopies unique to Mischka (median: 96.75%) vs. those found in either the fox or the dhole (median: 95.72%) tended to be slightly higher (p-value: 6.02*10^-44^) **(Supplementary Figure 6).**

To estimate the rate of retrocopy insertions, we first focus on the Mischka genome. We estimated the number of generations since genomic divergence from the mCanLor1.2 wolf genome. Excluding duplicated segments and short tandem repeats, there are 4,219,105 autosomal single nucleotide differences between the Mischka and mCanLor1.2 genome assemblies. Using the wolf mutation rate of 4.5 * 10^-9^ per generation [59], and with an analyzed genome size of 2,022,154,146 bp, we calculated that 231,827 generations have elapsed since genome divergence between Mischka and mCanLor1.2. By comparing the retrocopies between Mischka and mCanLor1.2, we identified 48 retrocopies that are unique to Mischka, and 79 that are unique to mCanlor1.2. Retrocopies that were present uniquely to Mischka had a much greater sequence similarity to their parent gene (mean: 99.92%) vs. those shared in the mCanLor1.2 assembly (mean: 96.14%) (p-value: 9.396*10^-29^) **(Supplementary Figure 6)**. By dividing the average amount of unique retrocopies over the number of generations, we estimate that the rate of retrocopy insertion as 2.74 * 10^-4^ per generation, or approximately 1 in 3,650 births, a higher rate than expected. Insertion rates estimated by comparing multiple canine assemblies, including Mischka, to the mCanLor1.2 wolf genome range from 2.74 * 10^-4^ per generation to 3.1 * 10^-4^ per generation, or from 1 in 3226 to 1 in 3650 (avg: 1/3514 births) **(Table 3)**. When considering a calibration based on the lower-bound and higher-bound wolf SNV mutation rate, which was defined as a range of 2.6 × 10^−9^ to 7.1 × 10^−9^ [59], we estimate the range of novel insertion in Mischka is 1.58 * 10^-4^ per generation to 4.32 * 10^-4^ per generation (1/2,315-1/6,329 births).

**Table 3.** Rate of Retrocopy Formation in Canine Assemblies. This table describes the five assemblies retained for retrocopy comparison against mCanLor1.2, the Greenland Wolf Assembly. Column 1: Assembly Name. Column 2 contains the total sum of retrocopies between the assembly and mCanLor1.2. Columns 3 and 4 show how many retrocopies are unique to the assembly and to mCanLor1.2 respectively. Column 5 shows the number of generations elapsed since divergence between the assembly in mCanLor1.2. Column 6 shows the final determined insertion rate, calculated by dividing the average between Columns 3 and 4 over the generations in Column 5, shown as an insertion rate of X * 10^-4^, and presented as 1 in n births. Column 7 shows the range of generations elapsed, based on the upper and lower bounds of the wolf mutation rate. Column 8 shows the range of insertion rate, based on the generation range in Column 7.

| **Assembly Name** | **Total Retrocopies Between Assemblies** | **Retrocopies Unique to Assembly** | **Retrocopies Unique to mCanLor1.2** | **Estimated Generations Since Divergence** | **Insertion Rate \* 10<sup>-4</sup> (1 in N births)** | **Range of Generations** | **Range of Insertion Rate \* 10<sup>-4</sup> (1 in N births)** |
| --- | --- | --- | --- | --- | --- | --- | --- |
| Mischka | 3,489 | 48 | 79 | 231,827 | 2.74 (3,650) | 146,932-401,239 | 1.58-4.32 (2,315-6,329) |
| China | 3,513 | 48 | 92 | 226,008 | 3.1 (3,226) | 143,245-391,168 | 1.79-4.89 (2,045-5,587) |
| Nala | 3,503 | 52 | 79 | 235,551 | 2.78 (3,597) | 149,293-407,685 | 1.61-4.39 (2,278-6,211) |
| Sandy | 3,510 | 54 | 79 | 234,571 | 2.83 (3,534) | 148,672-405,989 | 1.64-4.47 (2,237-6,098) |
| Zoey | 3,487 | 47 | 78 | 224,470 | 2.78 (3,597) | 142,270-388,507 | 1.61-4.39 (2,278-6,211) |

## Discussion

We analyzed the duplication content of nine recent canine assemblies using two methods: genome assembly self-alignment and read-depth analysis, identifying 1.74-7.05% of each genome as duplicated. We found that duplications were enriched in the first and last megabases of canine autosomes. Since canine autosomes are acrocentric [54], this suggests an enrichment of duplications in pericentromeric and subtelomeric regions, as found in other species [60].

The assembly self-alignment and read-depth approaches identified a range of duplication content in each assembly, ranging from 17.27 to 109.78 Mb by self-alignment and 41.17-166.47 Mb by read-depth. Overlap analysis shows a generally poor level of agreement, with most duplicated base pairs only identified by one of the methods. Thus, each of the assemblies offers an incomplete representation of the true duplication content in each sample. The assemblies differ in how they represent partially resolved duplications. For example, in the Mischka assembly, unresolved duplications were often retained as separate unplaced contigs that were not assigned to a chromosomal location. Other assemblies report few or no duplications on unplaced contigs. This difference may have implications in downstream analyses that utilize these assemblies.

Our initial analysis of duplications in the Mischka assembly revealed the presence of many short, high-recurrence duplications. Further investigation showed that these segments corresponded to multiple related retrogene insertions. To explore this further, we annotated 1,263-1,316 retrocopies across all assemblies, and found that a large majority displayed the hallmarks of retrotransposition such as poly(A) tails and target site duplications. This includes 49-66 retrogenes per assembly that appear to maintain intact parental ORFs. Whether these intact retrogenes are expressed or have any functional impact requires further investigation. We also note that our definition of retrogene boundaries is based on the existing annotation of genes in canines. This annotation of untranslated regions (UTRs) in these gene models is often incomplete, resulting in an underestimation of the size of retrogene insertions (**Figure 2B**), and underestimating the count of retrocopies with target-site duplications and/or poly(A) tails. We also find that 53% of retrocopies from Mischka are present in the dhole or fox assemblies, implying an origin prior to wolf speciation. However, it should be noted that the dhole and red fox assemblies are highly fragmented (n=29,680 and 82,424 contigs, respectively), limiting the ability to interpret sharing among the two genomes [57, 58].

Additionally, in the Mischka assembly, we find that 466 retrocopies are located within larger segmental duplications with over half of those retrocopies being present on unplaced contigs. This suggests a pattern where a retrocopy inserts into a locus, which is then subsequently duplicated, perhaps repeatedly, throughout the genome. A similar pattern of insertion-duplication has been described for retrogenes that may have functional effects in elephants and horses [61, 62].

We take advantage of retrocopy comparisons between dog and wolf assemblies to estimate an average retrogene insertion rate. Methods for estimating mutation rates from patterns of sequence differences can be divided into three broad categories [63]. First, rates can be estimated by sequencing trios and counting the number of new mutations observed in children that are absent in their parents. This estimate is the least affected by natural selection, as only lethal mutations should be unobserved in the offspring. However, a large number of individuals may need to be analyzed, particularly for mutation types that arise at a slow rate. This method is also extremely sensitive to false positive calls (in the children) and false negative calls (in the parents). Second, rates can be estimated from the distribution of variants identified in multiple individuals using population genetic models. The expected distribution of differences among individuals is a function of multiple population genetic parameters. If some parameters are known, an estimate of the mutation rate can be obtained. For example, if the effective population size is known, or can be estimated using other data, then the mutation rate can be estimated based on the distribution of segregating sites in the sample. Similarly, mutation rates can be estimated from differences between chromosome segments that are inherited identical-by-descent (IBD) if the number of generations separating the IBD segments can be determined.

Related methods have been applied over both recent and ancient time scales [64–67]. However, these methods may require models of how population sizes change over time, are sensitive to missing rare variants, and can be affected by natural selection. The third class of estimates takes a phylogenetic approach and estimates the mutation rate based on the number of differences that have occurred along two sampled lineages. The divergence time between sampled lineages can be estimated from species separation times inferred from the fossil record. However, for closely related groups, much of the lineage divergence time occurs as coalescence in the ancestral population, making estimates of the ancestral population size critical when a fossil calibration is used. As in population-based estimates, an alternative is to calibrate phylogenetic estimates based on a sequence divergence time estimated using a previously known rate for a different type of mutation.

In this study, we utilize phylogenetic comparisons between dog and wolf genomes to estimate the rate of retrocopy formation. We calibrate these estimates using a single nucleotide mutation rate inferred from sequencing a wolf pedigree. This mutation rate, 4.5*10^-9^/bp/gen, implies a divergence between assembled dog genomes and the mCanLor1.2 Greenland wolf of 231,827 generations, or 695,481 years, assuming 3 years per generation [68]. This deep divergence may seem inconsistent with the domestication of dogs in the past 15,000-30,000 thousand years [68–70]. Dogs are thought to have been domesticated from a now-extinct progenitor population of wolves somewhere in Asia and the ancestral wolf population may have had a large effective size. When correcting for differences in the assumed mutation rate, a previous Bayesian coalescent analysis of dog and wolf divergence estimated that the ancestral population size of new-world and old-world wolves was quite large, 143,000 individuals [71].

This implies a mean coalescence of two lineages in this ancestral population of 286,000 generations, which, along with the population separation time, is broadly consistent with our phylogenetically inferred estimate. We note that these estimates of population size may be biased by the presence of admixture from coyotes in some of new-world wolf samples used to create the estimate [71, 72] or by unmodeled population structure among ancestral wolves.

We note that the our initial annotation of retrocopies in each assembly (Table 2) applies conservative filters of alignment quality to reduce false positive annotations. However, this induces false negative calls across assemblies. For the rate estimate, we corrected for this by assessing the presence of retrocopies called in any assembly among all other assemblies through an alignment of the flanking sequences (Supplementary Figure 8). Although our estimate of 1/3,514 births is affected by several assumptions including the absence of selection and of a uniform rate, the estimate of the SNP mutation rate is likely the largest single source of uncertainty in our estimate. Utilizing the reported confidence interval of the SNP mutation rate (2.6*10^-9^ – 7.1*10^-9^) implies a retrogene insertion rate of between 1/2,226 and 1/6,075 births.

The inferred retrocopy insertion rate in canines is substantially higher than that found in humans; between humans and chimps, one new gene retrocopy insertion was estimated for every 6000 births, based on the 1000 Genomes Project [73]. However, this estimate is calibrated based on an assumed human effective population size of 10,000 which is largely consistent with an assumed human mutation rate of ∼2.5*10^-8^ bp/gen [74]. Adjusting this for the human mutation rate obtained from pedigree sequencing, ∼1.3 10^-8^ bp/gen [75], decreases the estimated rate of retrocopy insertion in humans to ∼1/11,500 births, or about 3-times lower than the canine estimate. However, it is noted that long-read data enables a higher retrocopy insertion detection rate [76] so this may be an underestimate of the true human insertion rate. Other observations are consistent with a high rate of retrogene formation in dogs. A recent survey suggests that polymorphic retrogene insertions are 13-fold more common in dogs than in humans [13]. The most well-studied retrogenes in dogs are derived from the *FGF4* gene which cause the short-leg phenotype found in some breeds such as corgis and beagles [12]. *FGF4* retrogenes have occurred multiple times independently, with 7 independent copies identified to date [13, 77]. Additionally, dogs have a high rate of SINE polymorphism [37, 78]. Both SINE and retrogene insertions require the activity of LINE-1 elements. In contrast to SINEs, which do not encode any proteins, LINE-1s are autonomous retroelements that encode proteins required for their own retrotransposition [79]. LINE-1 encoded proteins predominantly mobilize the RNA that encodes them through a process known as *cis-preference* [80]. However, one or more LINE-1 encoded proteins can rarely act to mobilize other RNAs such as SINEs or cellular mRNAs [81]. We speculate that the high rate of SINE and retrogene insertion found in dogs may result from a shared mechanism that involves a reduced level of *cis-preference* for the proteins encoded by the canine LINE-1 element.

## Materials and Methods

### Assemblies considered

Our analysis involves 9 published long-read canine assemblies [33–40]. For simplicity, we refer to them by name. Assembly accessions and other details are provided in **Table 1**.

### Genome assembly self-alignment

Genome assembly self-alignment was performed using BISER (Brisk Inference of Segmental duplication Evolutionary stRucture) (v1.2.3) which was developed to detect segmental duplications at low identity across multiple genome assemblies [16]. BISER defines a segmental duplication with the following criteria: below a certain error threshold based on shared sequence identity ε(approximately 15% for 75% shared sequence identity), longer than 1,000 bp, and paralog sequences may overlap up to a certain amount of base pairs, also defined by the error threshold. Before alignment, repetitive regions in the genome are soft-masked, prohibiting the initialization of alignments in low-complexity and highly-repetitive sequences, such as LINEs, SINEs, and simple repeats. Duplications involving the mitochondrial chromosome were removed from analysis since nuclear mitochondrial insertions (nuMTS) represent a distinct class of variation [82], and two categories of duplications were defined: one where the paralogous sequence between duplication copies is at least 90% identical, and a subset where the identity rate is at least 95%. These sets are further divided based on their placement on assembled chromosomes (the 38 canine autosomes and the X-chromosome) vs. unplaced contigs in each assembly. Because BISER identifies duplicated segments as pairs of sequences and duplication pairs often overlap each other, we used BEDTools merge (v2.30) to create a non-redundant set of duplicated intervals [83].

### Distribution of duplications along the genome

We analyzed the distribution of duplications located on the 38 canine autosomes and the X chromosome by permutation. The distribution of duplications was compared against 1000 random permutations using BEDTools shuffle (v2.30), limiting placement to the assembled chromosome sequence [83]. This was performed three times, on the self-alignment duplications, the read-depth duplications, and on the retrogene set to look for chromosomal enrichment.

### Locating highly duplicated segments

Short segments with a high recurrent duplication level were identified using the duplication pairs found by self-alignment located on the assembled chromosomes. All duplications smaller than 2.5 Kb were located, and their coordinates rounded to the nearest hundred (i.e., a coordinate of 2,349 would become 2,300, and a coordinate of 3,175 would become 3,200), to account for minor difference in alignment endpoints. Duplications were merged into segments, and segments that contained at least four duplications were labeled as highly recurrent.

### Read-depth analysis

Read-depth analysis was applied to Illumina data from the assembled individuals using fastCN [31]. First, a version of the assembly was created in which elements identified by RepeatMasker, tandem repeat finder [84], WindowMasker [85], and over-represented 50-mers were masked. Next, all placements within an edit distance of two for 36-bp segments of each Illumina read determined using mrsFAST (v3.4.1) [86]. The resulting depth profiles were tabulated, corrected for local GC content per fastCN, and converted to copy number estimates based on a set of control regions. For copy number normalization we excluded: reported deletions, duplications, and insertions [33], duplicated intervals from a preliminary self-alignment of the Mischka genome, duplicated segments identified by a preliminary fastCN analysis, and all chromosomes other than chr1-38. Copy number profiles were calculated based on the mean normalized depth tabulated in windows containing 1,000 unmasked positions.

Duplicated regions are defined as at least four consecutive windows whose copy numbers all exceed 2.5 and have a total combined size equal to or greater than 10 Kb. An overall copy number was determined by taking the median copy number of all windows within a duplicated region. Masking of the additional genomes was performed as described above. Normalization controls were converted from Mischka coordinates using minimap2 (v2.26) [87]. China was omitted from these analyses because, while it does have BGI short read sequencing, we opted to only utilize Illumina-based data. Nala was omitted due to extreme GC bias observed in the associated Illumina data **(Supplementary Figure 4)**.

### Comparison of duplications identified using self-alignment and read-depth approaches

To compare the results found by self-alignment and the results found by read-depth, both sets of duplications (BISER and read-depth) were filtered to retain only a) duplications on the assembled chromosomes; b) duplications greater than 15 Kb; c) duplications with at least 95% identity; d) duplications that are not located in the first megabase of any autosome. Analysis was limited to assembled chromosomes due to differences in the treatment of unplaced contigs between assemblies. The read-depth analysis has a mean window size of 3,822 bp, with four windows of an average size equaling approximately 15 kb. We utilized this metric to filter our read-depth and our self-alignment sets to match this new criterion. The first megabase of each autosome is excluded because, as a result of masking, there are no informative read-depth windows in these regions. Self-alignment duplications were intersected with 1 Kb unique sequence read-depth windows, and a self-alignment duplication was confirmed supported if the windows had a median copy number of at least 2.5. Read-depth duplications were counted as supported if they overlapped with any self-alignment duplication. The methods described above for the Mischka assembly were expanded to the eight other canine genome assemblies with control region coordinates using liftoff (v1.6.3) [87, 88]. In the gene analysis, read-depth windows are intersected with the protein-coding genes annotated in Mischka.

For the gene set of choice, we used a subset of 13,817 genes identified in Mischka, with the restriction that genes must be protein-coding, must be located on the 38 autosomes or the X chromosome, and that a gene must fully encompass at least 3 windows, where each window represents 1 Kb of non-repetitive DNA sequence used by fastCN. A table was built to observe copy number across the genome, where a row represents a canine sample and a column represented a pre-determined window in the Mischka coordinates, with entries being the copy number. Copy number for genes were determined as the median of all copy numbers in windows encompassed by a gene. A canine assembly is considered to have a duplicated gene if the copy number exceeds 2.5.

### Retrogene analysis

We searched for candidate retrocopies in the Mischka assembly based on the annotation of 19,972 protein coding genes used by the Dog10K consortium [14]. A single transcript was selected for each gene, with priority given to the longest isoforms and fewest number of exons. We removed any genes with an intron less than 50 bp, to account for many uninvestigated genes with single base pair introns, and any gene that only had a single exon.

We constructed cDNA sequences using SAMtools faidx (v1.13) to extract the sequence of each exon [89]. Using BLAT, cDNA sequences were mapped against the genome with repetitive 11mers filtered (*--ooc* option) (v.35) [90]. The resulting alignments were examined for hits that could be retrocopies. Alignments were removed if they were: less than 100 bp, overlapped the genomic coordinates of the parent gene, had less than 90% sequence identity with the parent gene, or do not cross an exon-exon boundary of the parent gene. The same analysis procedure was applied to the other eight genome assemblies after converting the genes annotated in Mischka to the coordinates of each assembly using liftoff based on minimap2 alignments (v2.26) [88, 91]. 1000 random permutations were conducted in the same manner described in the section *Distribution of duplications along the genome.* Due to paralogous genes, parent genes could not be resolved for a few retrocopies in each canine assembly.

### Conducting Gene Ontology (GO) Analysis

At several instances, large lists of genes were submitted to the Gene Ontology (GO) database in order to find functional links [44–46]. These were submitted to the GO Enrichment Analysis (Powered by Panther) under the ‘Biological Process’ tab to *Canis lupus familiaris*. A list of the genes submitted and their results can be found in **Supplementary Table 9**. We utilized the 2024-01-17 release (10.5281/zenodo.10536401).

### Examination of retrogene hallmarks

Retrogene containing loci from all nine canines were explored for presence of retrogene hallmarks: target-site duplications (TSDs), poly(A) tails, and LINE-1 endonuclease cleavage sites. Before detection proceeded, loci within one hundred base pairs of reference gaps were removed with BEDTools (v2.26.0) window with parameters *-w 100 and -u* [83]. Additionally, any loci within the first or last 100 base pairs of each chromosome as well as loci which were comprised of multiple segments did not undergo hallmark detection. For each locus, a symmetrical number of base pairs were extracted upstream and downstream of the retrogene with SAMtools faidx (v1.13) [89].

To detect TSDs, the upstream and downstream fasta files were aligned to each other using the water function within the EMBOSS module (v6.6.0) [92]. Water is a modified version of a Smith-Waterman local alignment and reports the highest scoring alignment only [93]. The scoring matrix used was a custom matrix which used +2 for all matches and -6 for all mismatches (-1000 for any alignment containing “N”). Gap opening and gap extension penalties were both set to 10. Alignment output containing “N” as a base pair were discarded, as were final alignments shorter than 5 base pairs in length. If alignments started within 5bp of the start of the upstream query or 5bp of the end of the downstream query, the flanking distance was increased by 5 and the alignment was repeated.

Poly(A) detection followed TSD detection. The original flanking distance worth of base pairs were again extracted beyond the 3’ end of the retrogene with SAMtools faidx (v1.13).

From there, strings of A or T, depending on retrogene orientation, were detected within the region. Poly(A)s were not extended into previously detected TSDs. If multiple putative poly(A) tails were detected, the poly(A) closest to the TSD was reported if one was present. If no TSD was detected, the Poly(A) closest to the retrogene was reported.

To assess if insertions possess a LINE-1 endonuclease cleavage site, loci which possess a detected TSD as determined above had the first five base pairs of TSD sequence extracted along with the two base pairs before the TSD using SAMtools faidx (v1.13). If the retrocopy is in the same orientation as the chromosome, the sequence is reverse complimented so that all cleavage site sequences are reported on the minus strand. Cleavage sites were then aggregated on a sample-by-sample basis and plotted using matplotlib [94] and logomaker (v0.8) [95] .

Various extract distances were trialed ranging from 10 base pairs to 2000 base pairs with 60 base pairs being chosen to maximize concordance between TSDs and poly(A) (concordance defined as the number of loci in which the TSD and poly(A) are within 5 bases of each other).

Seven of the nine samples had a maximum concordance with flanks of 50bp, 60bp, 70bp, 80bp, or 90bp, while the other two had a maximum concordance with flanks of 200bp. Though this method is limited in its ability to detect TSDs/poly(A) combinations that are more than 60bp combined, as well as hallmarks present in retrocopies containing unannotated UTR sequence, these limitations are thought to be minor as the vast majority of retrogenes contained at least one hallmark.

To assess fullORF loci, the list of retrocopies with full open reading frames was intersected with the loci which had their hallmarks identified (with 60bp of flanking distance) in the previous step using bedtools intersect (v2.26).

Polymorphic loci were investigated for an alternate pattern of TSDs or endonuclease cleavage sites. For each locus, an additional 5kb of flanking sequence of a retrocopy (or lack thereof, referred to as the “empty site”) was extracted using samtools faidx (v1.13). At each locus, canines with starting coordinates longer than 20,001bp were excluded from the analyses, as well as loci with multiple parent gene candidates. The extracted sequences were aligned to one another using age -both (v0.4) [97]. If an insertion was detected, the inserted sequence was used as a query to blat against the cDNA of the retrocopy of interest (v35). Blat commands followed as such: “blat -minIdentity=85 extracted_insertion_sequence cDNA_file blat_output_file”. The corresponding longest blat hit must be greater than 75% of the length of the insertion. Individual canine comparisons underwent separate processing if the blat alignment was of insufficient length, no insertion longer than 20bp was detected, or there was an insertion in both the query and the reference. These hits were not investigated in downstream steps. For remaining comparisons, AGE also detects locations of TSDs and is more stringent than our previous method, with detected TSDs being exactly at the boundary of the detected insertion and requires perfect sequence identity between upstream and downstream TSDs. For loci that contained a TSD, an endonuclease cleavage site was determined by extracting the last two bases before the TSD and the first 5 bp of the TSD using samtools faidx (v1.13). Cleavage sites were reverse complimented if the orientation of the retrocopy was forward. Logo plots and histograms of this dataset were generated using logomaker(v0.8) and matplotlib [95]. At each locus, only one confirmed filled site was processed. Priority was given to canines with at least 5bp of TSD sequence in the following order: mCanLor, then Mischka, Sandy, Nala, China, Zoey.

### Estimating the insertion rate of retrocopies

We selected contiguous genome assemblies from diverse dog samples to search for the insertion rate of retrocopies, including Mischka, Nala, China, Sandy, and Zoey relative to the mCanLor1.2 assembly. We discounted all retrocopies that overlap a read-depth segmental duplication. For each canine retrocopy set, pairs of 500 bp flanks were extracted from each retrocopy and aligned to mCanLor1.2 using minimap2 (v2.26) with the -c and -x asm5 options [87]. Retrocopies whose flanks did not map were discounted entirely from the analysis, due to an inability to determine presence of sharing between samples. For all retrocopies, if a retrocopy’s presence across all six assemblies had not yet been determined in the initial mapping, both the flanks and the intervening sequence were derived from an assembly that did have the sequence (in order, priority was given to mCanLor, then Mischka, Sandy, Nala, China, and Zoey in that order) and mapped to all dogs to determine whether or not the retrocopy was truly present. If the flanks were present and the intervening sequence was also present, the retrocopy was marked as shared. If only the flanks were present, the retrocopy was marked as absent in that dog, and if flanks did not map, the entire retrocopy was discounted, once again due to inability to determine sharing. The remaining retrocopies were totaled for presence or absence in each assembly, and converted into an upset plot using UpSet (v.0.9.0) [96].

Then, a retrocopy insertion rate was estimated based on a comparison between each assembly and mCanLor1.2. For each independent comparison, a retrocopy was considered as long as it could be determined whether or not it was present in mCanLor1.2 and the compared assembly, regardless of whether it failed to map in another assembly. This allowed for determination of retrocopies unique to mCanLor1.2 vs. a comparison assembly.

To identify sequence identity of retrocopies to the parent gene, retrocopy sequences were compared against the parent gene cDNA using BLAT (v.35) [90]. Sequence identity was determined as number of matches over number of matches plus mismatches. Retrocopies were then divided into categories for comparison: one group with fullORF retrocopies vs. those that were not; one group with retrocopies that were also found in either the dhole or the fox vs. those that were not; and one group with retrocopies unique to the Mischka assembly vs. those that were also found in the mCanLor1.2. assembly.

To further estimate rates of differences between genome assemblies, we identified single nucleotide differences among assemblies. Autosomal assembly size was determined by called regions between an assembly and mCanLor, as determined by minimap2 (v2.20) [87], with the additional flags of *-c --cs*, outputting differences in *PAF* format using the paftools.js call command. Regions are filtered to remove duplicated segments identified by read-depth, duplications identified by self-alignment, and simple tandem repeats identified by Tandem Repeat Finder [84]. SNPs called by minimap2 were then intersected with filtered regions. The resulting SNPs were used to estimate the number of generations that had passed since divergence between mCanLor1.2 and the other genome assemblies. We utilize the wolf SNP mutation rate estimated by Koch et al. of 4.5 * 10^-9^ per generation [59] and the size of the analyzed genome after filtering to determine generations passed as:

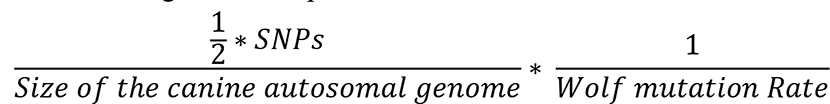

This formula calculates the approximate number of generations since divergence. We also calculated a range for the generations based on the range of the wolf mutation rate, which was 2.6 × 10^−9^ and 7.1 × 10^−9^. We divide the average number of unique retrocopies detected between a given canine assembly and mCanLor1.2 and divide by the estimated generational since divergence to get an estimate for the rate of retrocopy insertion, and additionally calculated a range of rate of insertion based on the generation range [25, 97, 98].

### Locating ancestral retrocopies

We selected 3,658 retrocopies from the Mischka assembly to search for in the red fox (*Vulpes vulpes,* VulVul2.2) and dhole (*Cuon alpinus,* GWHAAAC00000000) genome assemblies. It should be noted that both genome assemblies are composed of contigs, as opposed to chromosomes. We extracted 1 Kb flanks from all the retrocopies with samtools faidx (v1.13) and used minimap2 (v2.26) with the -c and -x asm20 options to align these flanks to the fox or dhole genome. We filtered out any retrocopies whose flank pairs either had at least one member missing, or whose flank pairs mapped to different contigs. We then took the intervening sequence between the flanks and removed any that were not at least 75% of the size of the original retrocopy in Mischka. With the remaining set, we extracted the intervening sequence with samtools faidx from their respective genomes and used minimap2 with the same options to map the putative retrocopies back to the Mischka assembly. If a candidate retrocopy from the fox or dhole did not overlap the original location in Mischka, it was discounted from the analysis.

## Acknowledgements

We thank the numerous authors of the genome assemblies used in this analysis for publicly providing their data for the investigation. In addition, we thank the editor and anonymous reviewers for their helpful comments.

## Author Contributions

AKN and JMK conceived the study. AKN carried out the data-gathering and primary analyses, and MSB analyzed retrocopies for retrotransposition hallmarks. MSB and JMK advised on the methodology and interpretation of results. AKN drafted the manuscript, and all authors reviewed and edited the final manuscript.

## Data Availability

Final counts and coordinates of each set of items (self-alignment duplications, read-depth duplications, genes in read-depth duplications, and retrogenes with hallmarks) are located in the provided supplementary files. Relevant and custom code is available on GitHub at https://github.com/antnguye/dups_and_retrogenes_in_canine_assemblies. Hallmark analysis is available on GitHub at https://github.com/mblacksmith1996/Hallmark_Detection_Python_Scripts. A UCSC genome browser TrackHub depicting duplications identified in the Mischka and mCanLor1.2 assemblies is available at https://github.com/KiddLab/dog-segdup-tracks.

## Supplementary Figures

**Supplementary Figure 1.**
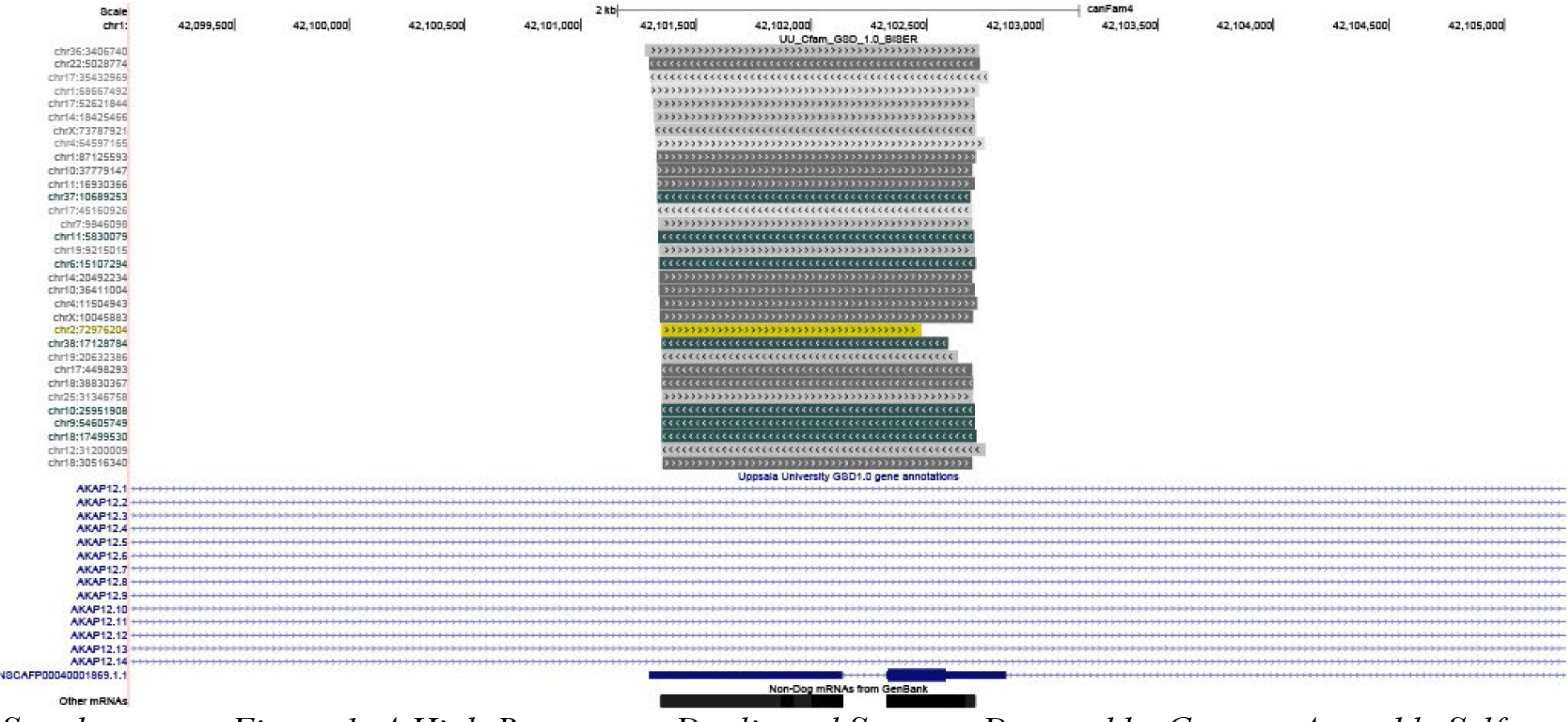
A High-Recurrence Duplicated Segment Detected by Genome Assembly Self- Alignment. This screenshot of UCSC Genome Browser depicts the region chr1:42,099,050-42,105,269 in Mischka, a segment of the genome located in an intron of the gene AKAP12. The green and gray bars depict duplicated intervals detected by genome assembly self-alignment (BISER, top), whereas the blue lines with arrows indicate genes annotated in the Mischka assembly. The final track represents non-canine mRNAs from GenBank. Each gray/green bar has a location where the duplicated sequence is found elsewhere as indicated at locations given on the left at homology levels between 90 and 98%, with the singular yellow bar indicating an even greater homology between 98% and 99%. The parental retrogene for this duplicated sequence is HMGN2, a common parent gene.

**Supplementary Figure 2.**
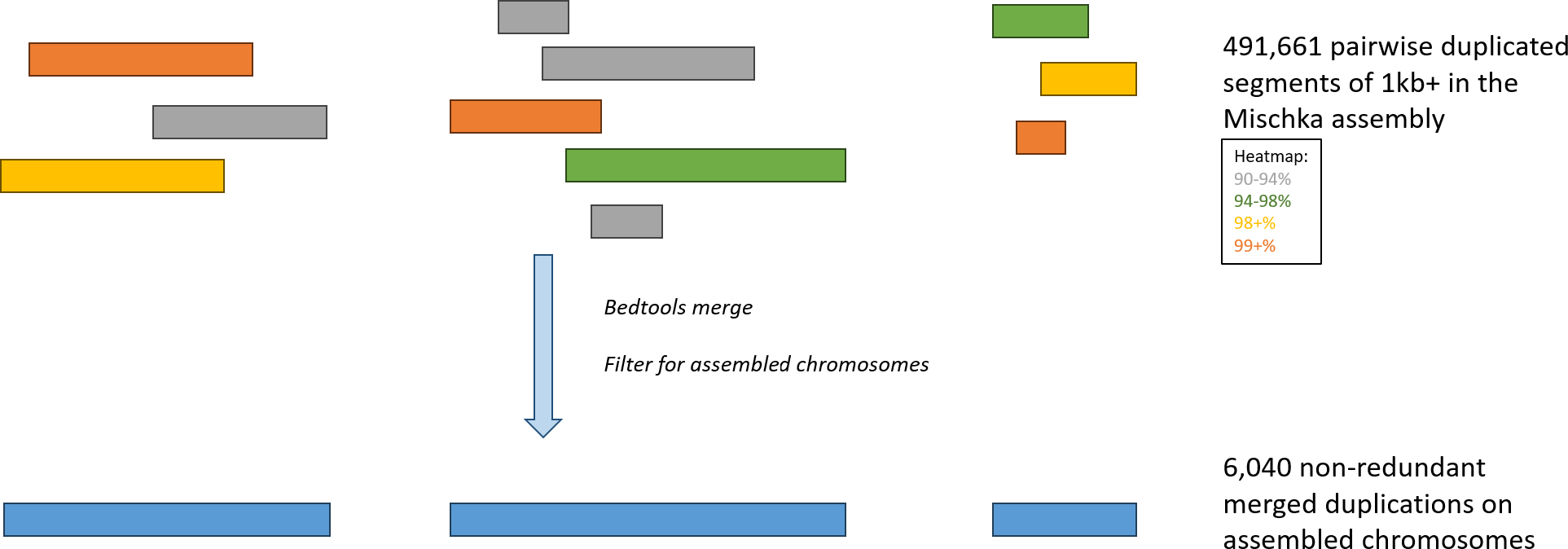
Merging Pairwise Duplications Identified by Genome Self Alignment. In the genome assembly self-alignment analysis, we identify pairwise duplicated segments 1kb in size with greater than 90% sequence similarity (represented in color). These duplicated segments appear across all chromosomes present in canine assemblies, including on unplaced contigs. Because these duplications are pairwise, a single locus can be duplicated multiple times and be present in multiple pairwise alignments with varying levels of sequence similarity (depicted in color). When merging, bedtools merge (v2.30) was used to combine overlapping segments together, regardless of sequence similarity We also removed duplications present on non-assembled chromosomes at this stage (chrM and unplaced contigs), since the treatment of unplaced contigs varies among the assemblies we studied.

**Supplementary Figure 3.**
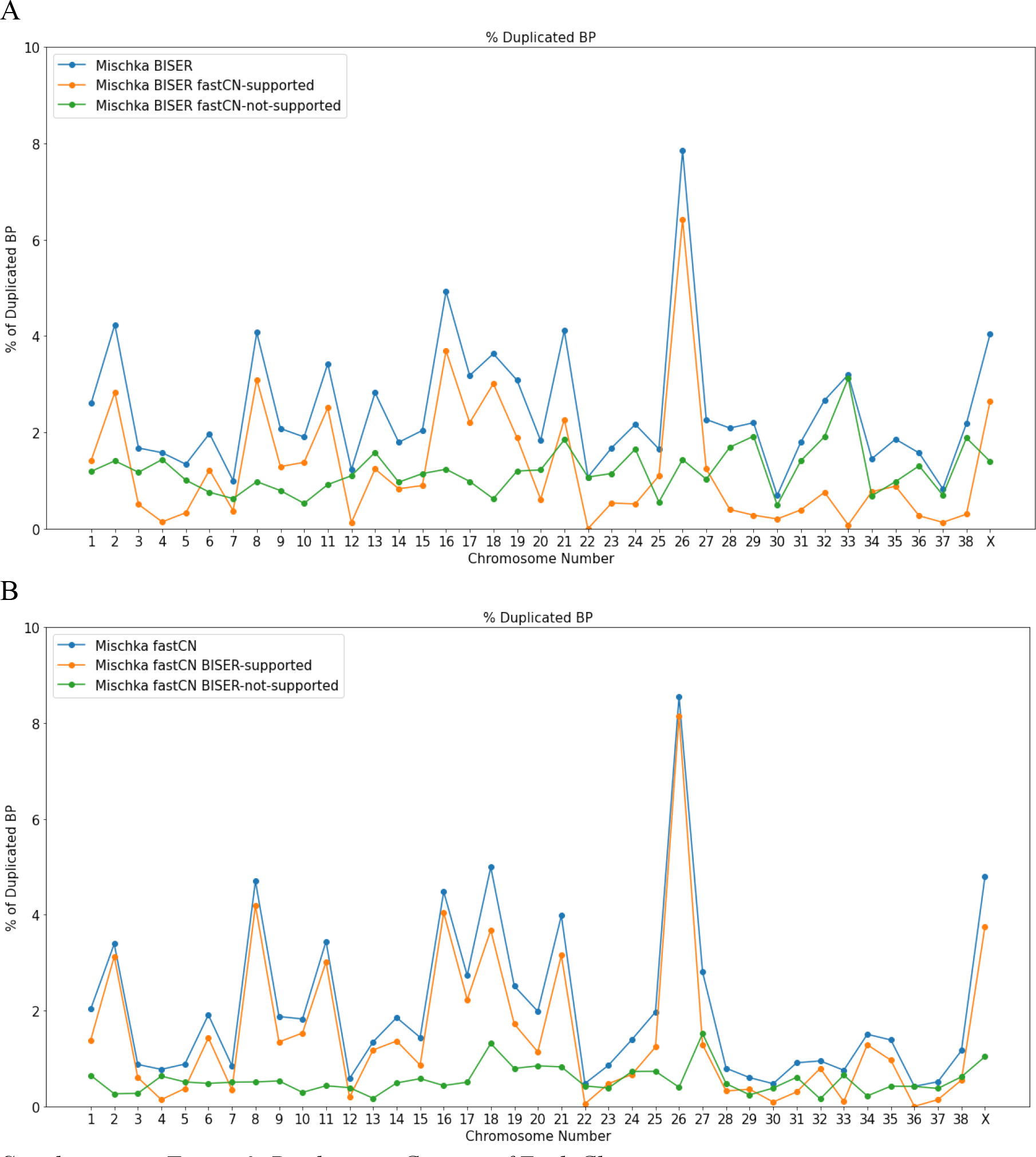
Duplication Content of Each Chromosome. The X-axis represents each assembled chromosome, and the Y-axis represents what fraction of that chromosome is duplicated as a percentage. (A) Blue represents duplications found by genome assembly self-alignment (BISER). Orange is BISER duplications supported by read-depth, and green is unsupported. (B) Blue represents duplications detected by read-depth. Orange shows support from BISER, and green is unsupported.

**Supplementary Figure 4.**
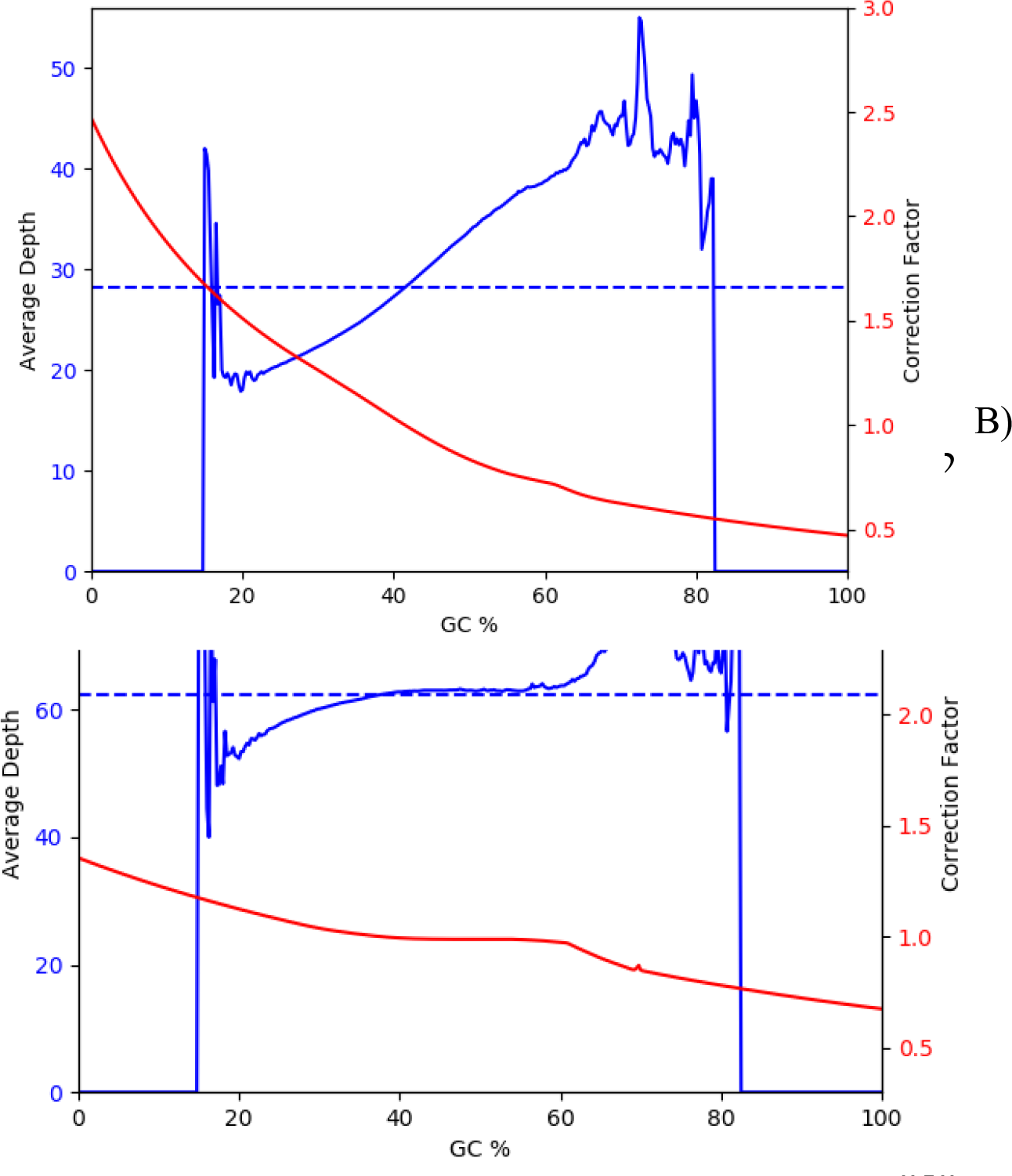
GC Coverage Profiles. Plots of coverage by local GC content are produced by fastCN. The dashed blue line shows mean read-depth. The solid blue line shows the coverage as a function of local GC content. The red line shows the correction factor calculated to normalize coverage based on GC content. (A) Depicted is the curve for Nala, which shows a skewed coverage across different GC levels, suggesting bias in the Illumina data. (B) Depicted is the curve for Mischka, showcasing a more uniform coverage profile across GC levels.

**Supplementary Figure 5.**
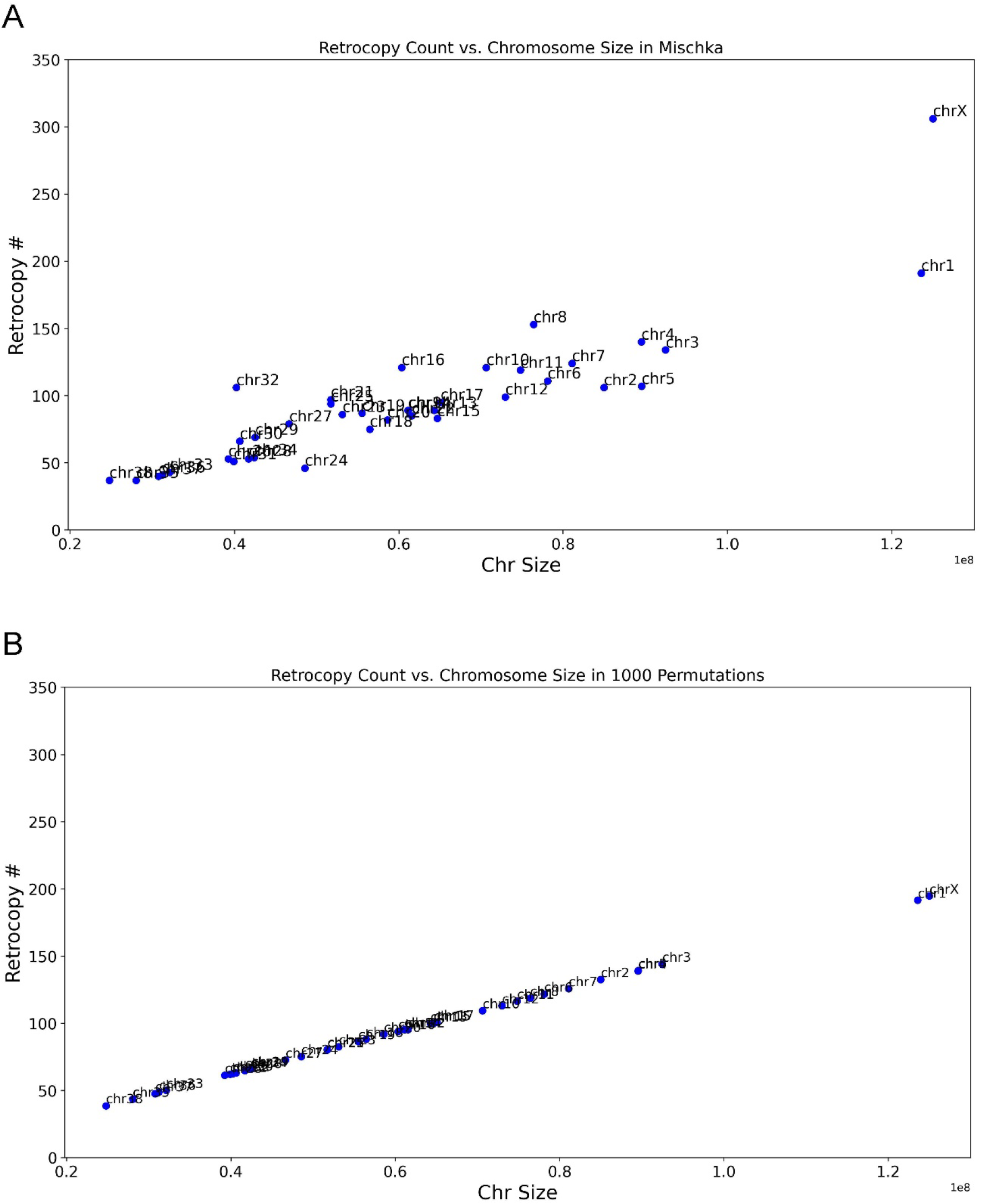
Retrocopy Count vs. Chromosome Size. A) Depiction of the number of retrocopies on each of the assembled chromosomes (chr1-38 + X) found in the Mischka assembly vs. the length of each chromosome. ChrX has 228 total retrocopies. B) Depiction of the number of retrocopies on each of the assembled chromosomes found in 1000 random permutations vs. the length of each chromosome. ChrX has a mean of 146 retrocopies and a max of 188 retrocopies. Mischka contains a 1.18-fold increase of detected retrogenes on chrX.

**Supplementary Figure 6.**
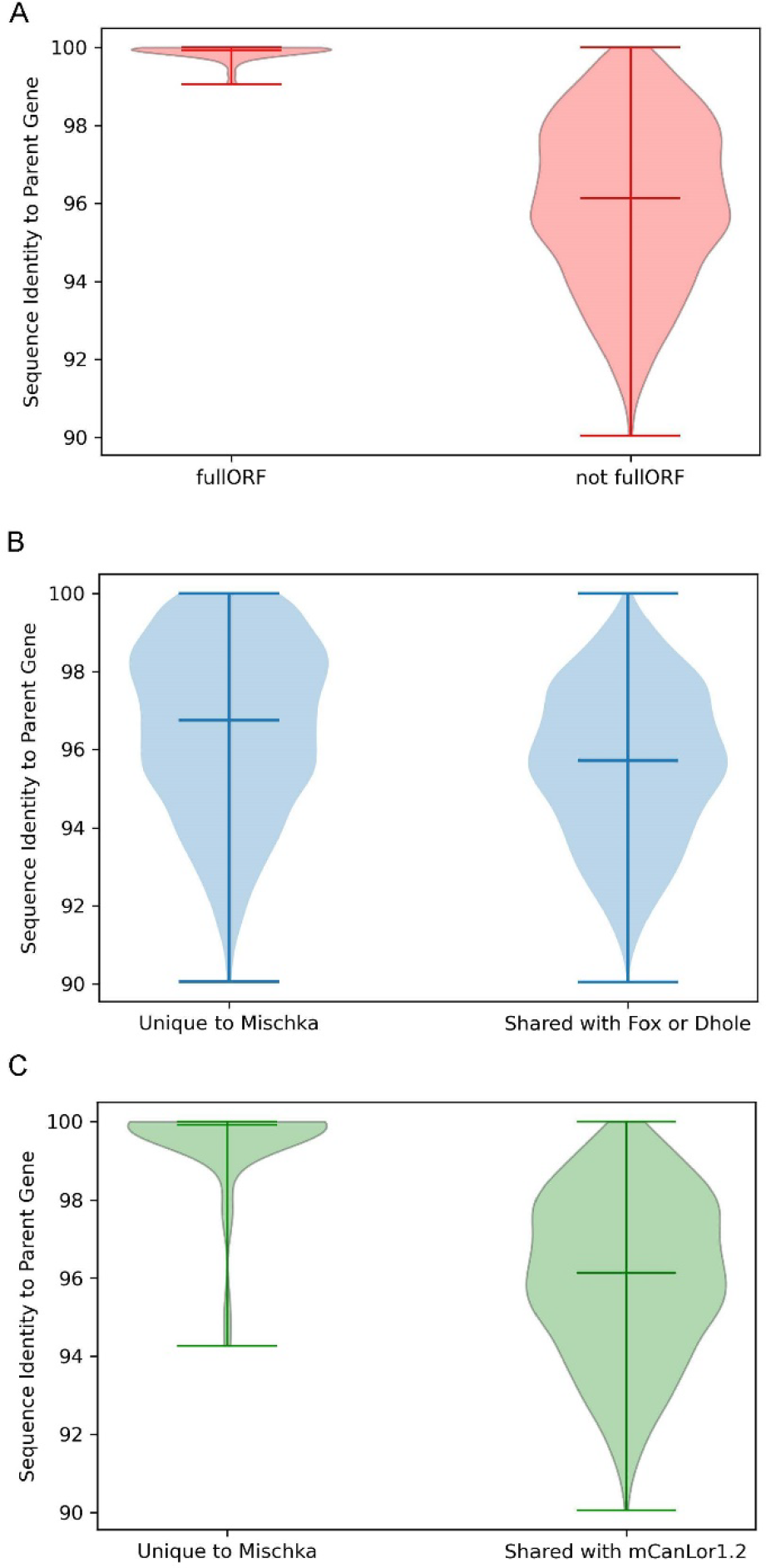
Sequence Similarity Between Mischka Retrocopies and Parent Gene This violin plot depicts three categories of comparison of sequence similarity between retrocopies found in the Mischka assembly and the parent gene. A) Sequence similarity of retrocopies between those that retain the parental open-reading frame (Mean: 99.86%; Median: 99.93%) vs. those that do not (Mean: 96.04%; Median: 96.14%). P-value: 3.81*10^-48^. B) Sequence similarity of retrocopies unique to Mischka (Mean: 96.57%; Median: 96.75%) vs. those found in either the dhole or the fox assemblies (Mean: 95.62%; Median: 95.72%) P-value: 6.01*10^-44^. C) Sequence similarity of retrocopies unique to Mischka (Mean: 99.54%; Median: 99.92%) vs. those found in the mCanLor1.2 assembly (Mean: 96.04%; Median: 96.14%). P- value: 9.396*10^-29^.

**Supplementary Figure 7.**
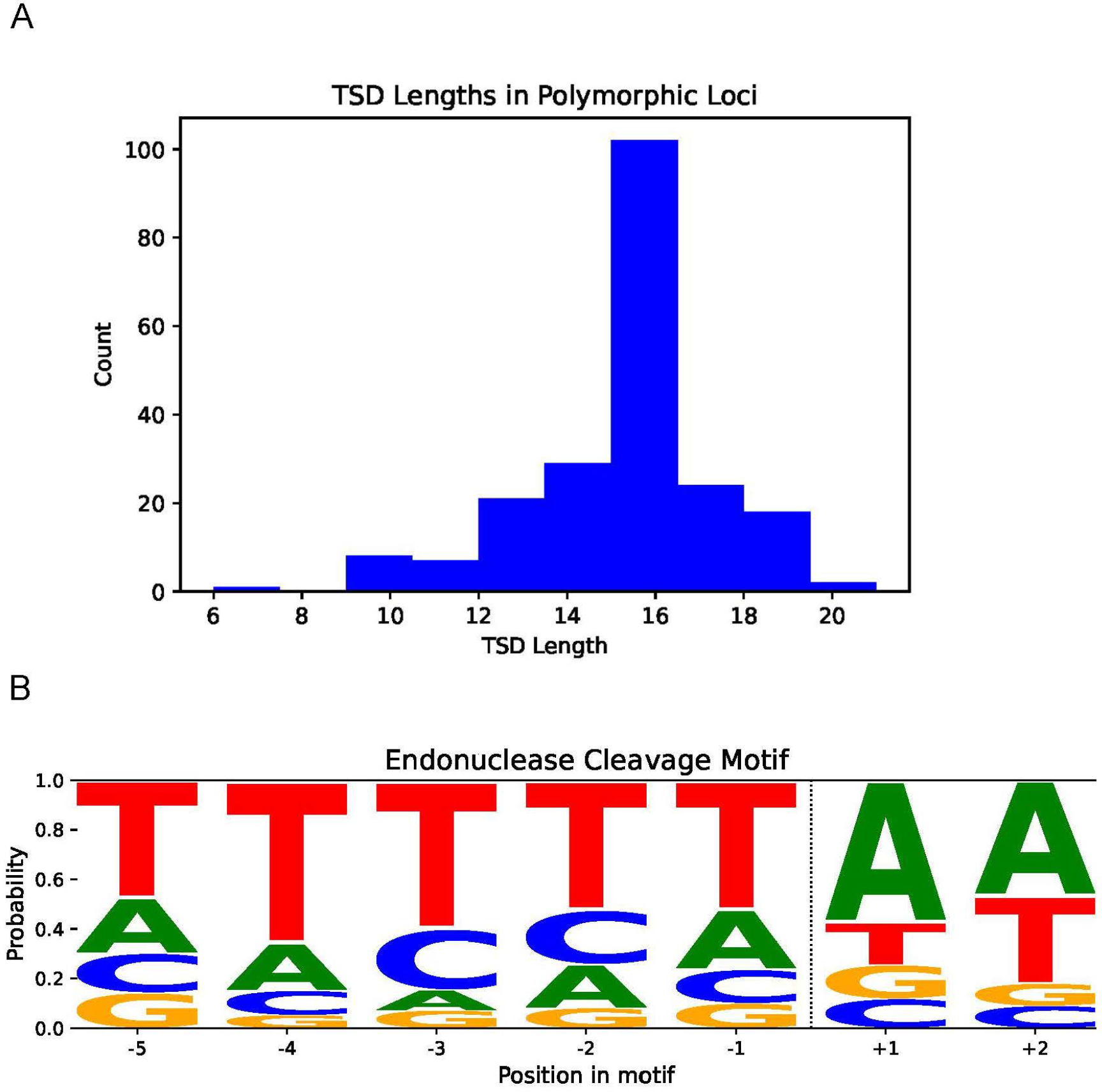
Hallmark Analysis of Polymorphic Retrocopies. An analysis of the polymorphic retrocopies (n=226 resolved loci). A) Detected TSD lengths for all polymorphic retrocopy containing loci with TSDs 5bp or longer (N=212). Only one canine is represented at each locus. Priority is given as to the first canine with an identified TSD as follows mCanLor, then Mischka, Sandy, Nala, China, and Zoey. The average TSD length across the dataset is 15.1 bp. B) The endonuclease cleavage motif for polymorphic loci containing 5bp TSDs, which follows the expected sequence of TTTTT/AA.

**Supplementary Figure 8.**
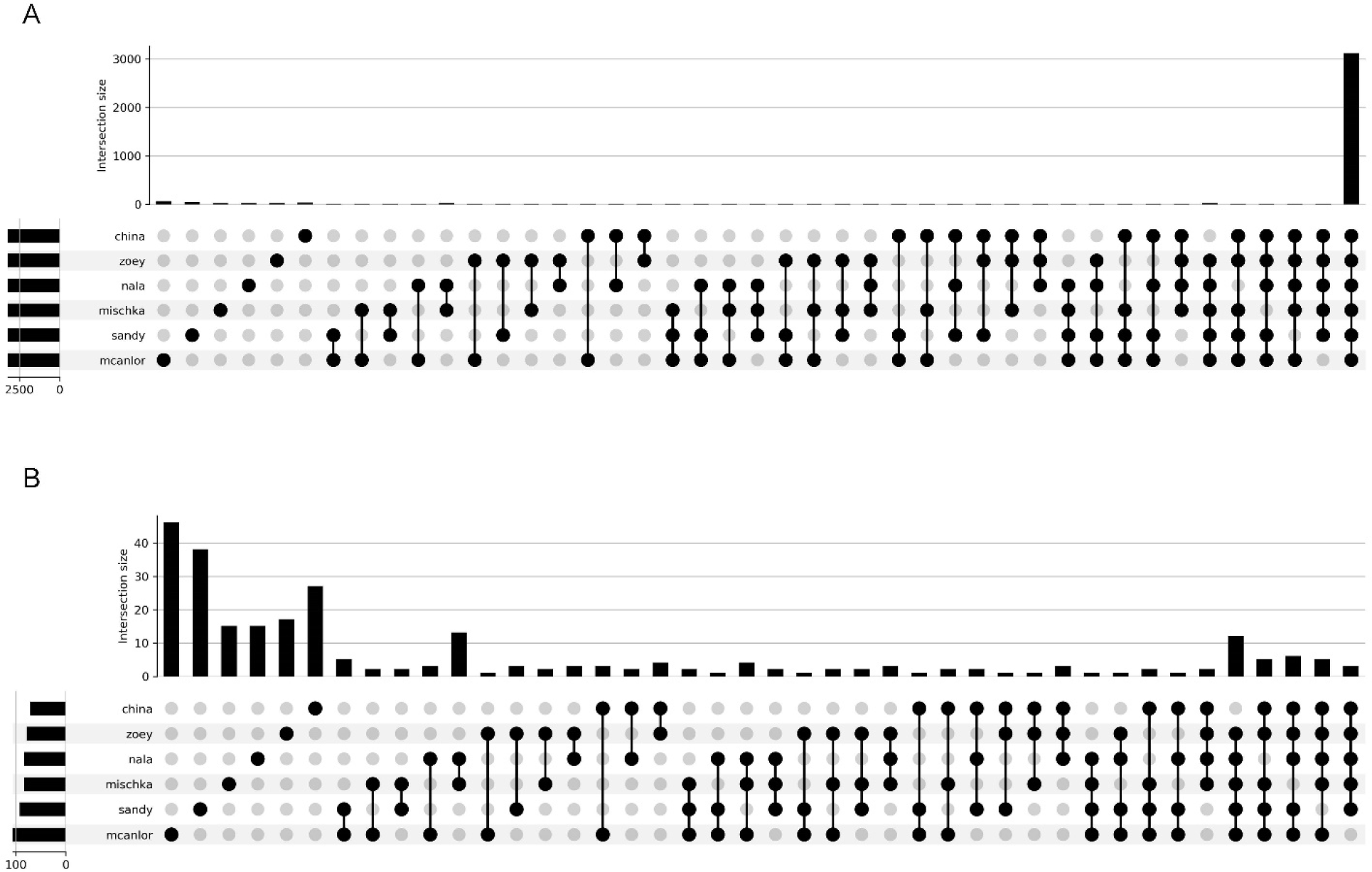
Retrocopies Shared Across Canine Assemblies. This upset plot depicts 3,377 retrocopies that were found to be present in at least one assembly between Mischka, mCanLor1.2, Sandy, China, Nala, and Zoey. A) depicts all retrocopies grouped by presence in canine assemblies, and B) removes the category for shared between all six assemblies. Of the retrocopies depicted, 3,111 are shared between all six assemblies. The second largest category is mCanLor1.2-specific retrocopies (n=56). The remaining categories have between 1 and 38 retrocopies.

## Supplementary Tables

**Supplementary Table 1.** All Duplications found by Genome Assembly Self-Alignment. Column 1 indicates the canine assembly’s name. Column 2 indicates the number of merged duplications detected by genome assembly self-alignment. Column 3 indicates the total amount of base pairs, in mBP, occupied by column 2, along with the percentage of the genome represented. Columns 4 and 5 are the same as 2 and 3, but filtered to only include duplicated intervals greater than 10 kB in size, and higher than 95% sequence identity. Column 6 is the mBP of self-alignment duplications located on unplaced contigs, and the percentage of base pairs within contigs that are considered duplicated per assembly.

| Name | Self-alignment Duplication Count | mBP occupied by duplications (% of the genome) | Self-Alignment Duplications filtered to 10kB and 95% Homology | mBP occupied by Self Alignment duplications filtered to 10kB and 95% (% of the genome) | mBP occupied by BISER duplications on unplaced contigs (% of the genome) |
| --- | --- | --- | --- | --- | --- |
| Mischka | 7980 | 97.68 (4.15%) | 3293 | 82.22 (3.49%) | 115.79 (90.15%) |
| mCanLor1.2 | 5256 | 134.54 (5.58%) | 781 | 109.78 (4.55%) | 15.78 (53.05%) |
| China | 5535 | 52.99 (2.27%) | 841 | 27.35 (1.17%) | 5.46 (34.73%) |
| Nala | 5887 | 111.84 (4.69%) | 1384 | 84.53 (3.55%) | 5.41 (23.58%) |
| Sandy | 5490 | 56.65 (2.42%) | 848 | 30.79 (1.32%) | 4.22 (33.31%) |
| Tasha | 5571 | 42.26 (1.83%) | 566 | 17.27 (0.75%) | 0.43 (20.14%) |
| Wags | 6993 | 94.4 (4.13%) | 2398 | 72.39 (3.17%) | 44.67 (36.98%) |
| Yella | 6751 | 103.47 (4.32%) | 1139 | 68.66 (2.87%) | N/A |
| Zoey | 6571 | 55.9 (2.4%) | 1494 | 31.36 (1.35%) | 11.73 (69.54%) |

**Supplementary Table 2.** All Duplications found by Read-depth. Duplications were defined as at least four consecutive windows found by fastCN to have a copy number of at least 2.5, with a size cutoff of 10 Kb. Column 1 contains canine name, column 2 contains the total number of duplications detected by read-depth, column 3 contains the total number of duplicated Mbp, and columns 4 and 5 mimics 2 and 3, but restricted to chr1-38 + X, referred to as the “assembled chromosomes”.

| Name | Read-depth Duplications | Duplicated Mbp (% of genome) | Read-depth Duplications on Assembled Chromosomes | Mbp on Assembled Chromosomes (% of assembled genome) |
| --- | --- | --- | --- | --- |
| Mischka | 1573 | 87.1873 (3.5%) | 1033 | 56.2099 (2.39%) |
| mCanLor1.2 | 2093 | 130.4306 (5.33%) | 2069 | 123.14 (5.11%) |
| Nala | 4709 | 152.5044 (6.34%) | 4706 | 152.3764 (6.39%) |
| Sandy | 1702 | 57.157 (2.43%) | 1681 | 55.1293 (2.36%) |
| Tasha | 990 | 42.5327 (1.84%) | 990 | 42.5327 (1.84%) |
| Wags | 1007 | 63.9808 (2.65%) | 688 | 45.5681 (1.99%) |
| Yella | 1060 | 98.3779 (4.11%) | 1059 | 98.3612 (4.11%) |
| Zoey | 929 | 48.9654 (2.09%) | 695 | 40.4794 (1.74%) |

**Supplementary Table 3.**
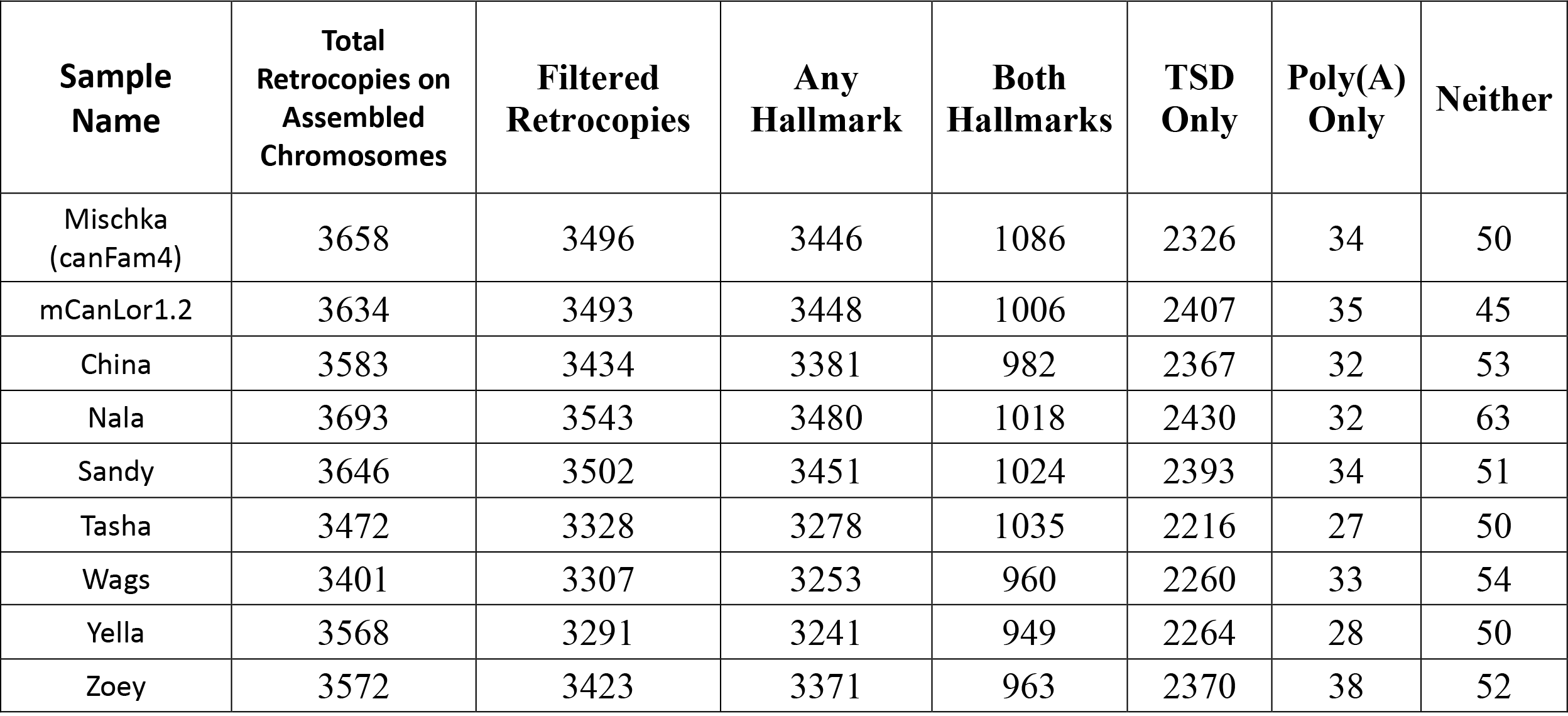
Hallmarks found in Retrocopies on Assembled Chromosomes. Total retrocopies found in Table 4 were filtered prior to hallmark analysis. Retrocopies were only considered for hallmark analysis if they were at least 100 bp away from either end of a chromosome, did not overlap any genome assembly gap, and were on chr1-38 + chrX. Hallmarks were only searched for within 60 bp outside of a retrocopy, with total number found in Column 3: Filtered Retrocopies. Target site duplications (TSDs) were required to be a minimum of 5 bp. Numbers correspond to the number of retrocopies that have the presence of either hallmark, both hallmarks, only target site duplications, only poly(A) tails, or no evidence of either hallmark.

**Supplementary Table 4.**
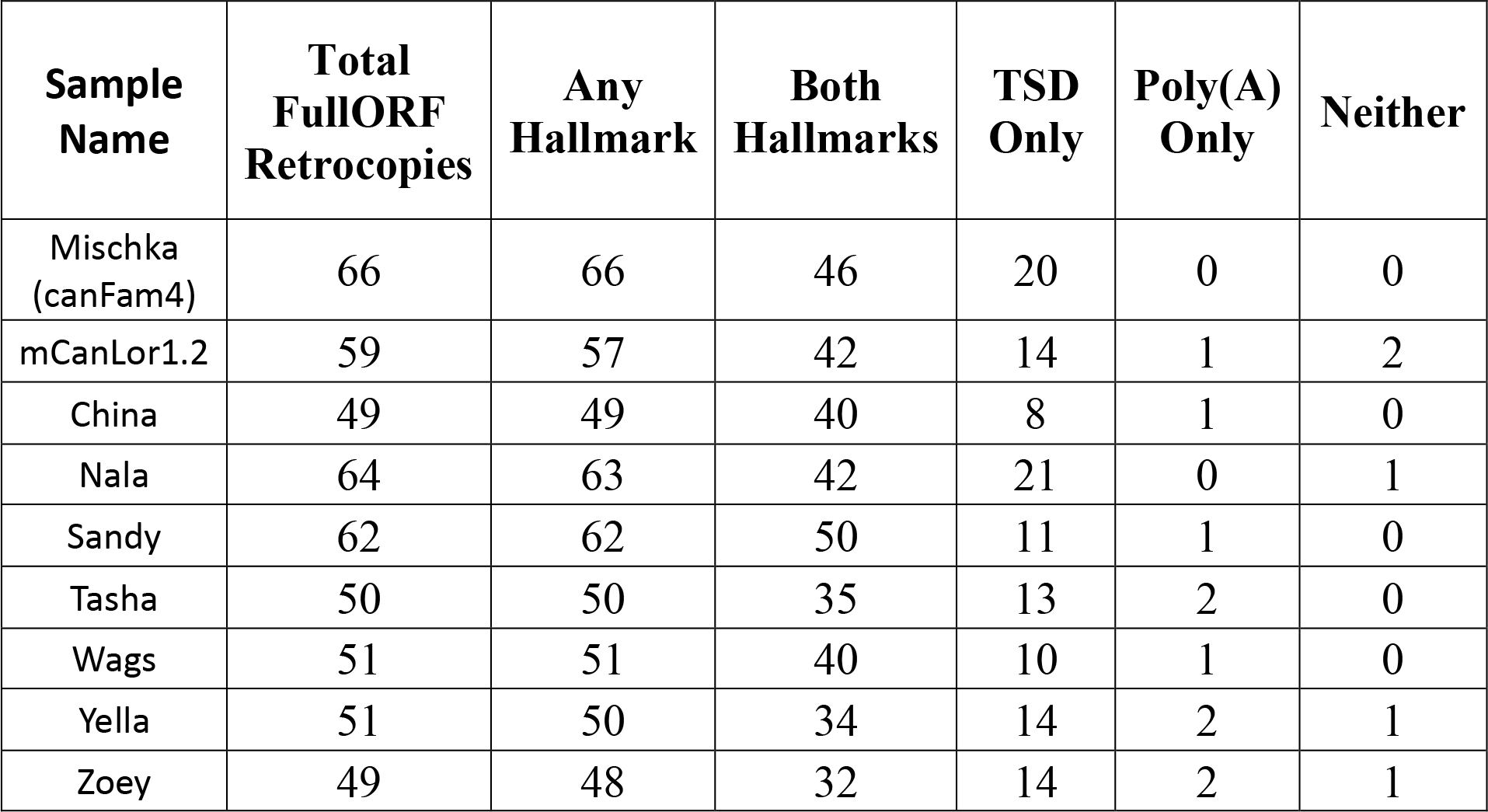
FullORF Retrocopy Hallmarks. This table mimics the same pattern as seen in Table 2, except the only retrocopies present in this table retain the full open reading frame as their parent gene. A majority of fullORF retrocopies have target-site duplications, with extremely rare occurrences without. 65-81% of fullORF retrocopies have evidence of both hallmarks.

| **Sample Name** | **Total FullORF Retrocopies** | **Any Hallmark** | **Both Hallmarks** | **TSD Only** | **Poly(A) Only** | **Neither** |
| --- | --- | --- | --- | --- | --- | --- |
| Mischka (canFam4) | 66 | 66 | 46 | 20 | 0 | 0 |
| mCanLor1.2 | 59 | 57 | 42 | 14 | 1 | 2 |
| China | 49 | 49 | 40 | 8 | 1 | 0 |
| Nala | 64 | 63 | 42 | 21 | 0 | 1 |
| Sandy | 62 | 62 | 50 | 11 | 1 | 0 |
| Tasha | 50 | 50 | 35 | 13 | 2 | 0 |
| Wags | 51 | 51 | 40 | 10 | 1 | 0 |
| Yella | 51 | 50 | 34 | 14 | 2 | 1 |
| Zoey | 49 | 48 | 32 | 14 | 2 | 1 |

## Supplementary Files

Table S5A. All Pairwise Self-Alignment Duplications Table S5B. All Self-Alignment Duplicated Intervals

Table S6. All Read-depth Segmental Duplications

Table S7. Duplicated Genes found by Read-depth

Table S8. All Retrocopies across All Assemblies

Table S9. GO Analysis of Duplicated and Parent Genes

Table S10. Shared Retrocopies Across Selected Canine Assemblies Table S11. Mischka Retrocopies Converted to Fox and Dhole

## References

1. Freeman JL, Perry GH, Feuk L, Redon R, McCarroll SA, Altshuler DM, Aburatani H, Jones KW, Tyler-Smith C, Hurles ME, et al: Copy number variation: new insights in genome diversity. Genome Res 2006, 16:949–961.

2. Conrad B, Antonarakis SE: Gene Duplication: A Drive for Phenotypic Diversity and Cause of Human Disease. Annual Review of Genomics and Human Genetics 2007, 8:17–35.

3. Dehal P, Boore JL: Two Rounds of Whole Genome Duplication in the Ancestral Vertebrate. PLOS Biology 2005, 3:e314.

4. Abdullaev ET, Umarova IR, Arndt PF: **Modelling segmental duplications in the human genome**. BMC Genomics 2021, 22:496.

5. Rody HVS, Baute GJ, Rieseberg LH, Oliveira LO: Both mechanism and age of duplications contribute to biased gene retention patterns in plants. BMC Genomics 2017, 18:46.

6. Payen C, Koszul R, Dujon B, Fischer G: Segmental Duplications Arise from Pol32-Dependent Repair of Broken Forks through Two Alternative Replication-Based Mechanisms. PLOS Genetics 2008, 4:e1000175.

7. Du RQ, Lu CC, Jiang ZW, Li SL, Ma RX, An HJ, Xu MF, An Y, Xia YK, Jin L, et al: Efficient typing of copy number variations in a segmental duplication-mediated rearrangement hotspot using multiplex competitive amplification. Journal of Human Genetics 2012, 57:545–551.

8. Esnault C, Maestre J, Heidmann T: Human LINE retrotransposons generate processed pseudogenes. Nat Genet 2000, 24:363–367.

9. Kaessmann H, Vinckenbosch N, Long MY: **RNA-based gene duplication: mechanistic and evolutionary insights**. Nature Reviews Genetics 2009, 10:19–31.

10. Casola C, Betran E: The Genomic Impact of Gene Retrocopies: What Have We Learned from Comparative Genomics, Population Genomics, and Transcriptomic Analyses? Genome Biol Evol 2017, 9:1351–1373.

11. Tutar Y: Pseudogenes. Comparative and Functional Genomics 2012.

12. Brown EA, Dickinson PJ, Mansour T, Sturges BK, Aguilar M, Young AE, Korff C, Lind J, Ettinger CL, Varon S, et al: FGF4 retrogene on CFA12 is responsible for chondrodystrophy and intervertebral disc disease in dogs. Proceedings of the National Academy of Sciences 2017, 114:11476–11481.

13. Batcher K, Varney S, York D, Blacksmith M, Kidd JM, Rebhun R, Dickinson P, Bannasch D: **Recent, full-length gene retrocopies are common in canids**. Genome Res 2022, 32:1602–1611.

14. Meadows JRS, Kidd JM, Wang GD, Parker HG, Schall PZ, Bianchi M, Christmas MJ, Bougiouri K, Buckley RM, Hitte C, et al: Genome sequencing of 2000 canids by the Dog10K consortium advances the understanding of demography, genome function and architecture. Genome Biol 2023, 24:187.

15. Bianchi CA, Marcellin-Little DJ, Dickinson PJ, Garcia TC, Li CF, Batcher K, Bannasch DL: FGF4L2 retrogene copy number is associated with intervertebral disc calcification and vertebral geometry in Nova Scotia Duck Tolling Retrievers. Am J Vet Res 2023, 84.

16. Išerić H, Alkan C, Hach F, Numanagić I: Fast characterization of segmental duplication structure in multiple genome assemblies. Algorithms for Molecular Biology 2022, 17:4.

17. Salzberg SL, Yorke JA: **Beware of mis-assembled genomes**. Bioinformatics 2005, 21:4320–4321.

18. Pan W, Lonardi S: Accurate detection of chimeric contigs via Bionano optical maps. Bioinformatics 2019, 35:1760–1762.

19. Hartasanchez DA, Braso-Vives M, Heredia-Genestar JM, Pybus M, Navarro A: Effect of Collapsed Duplications on Diversity Estimates: What to Expect. Genome Biol Evol 2018, 10:2899–2905.

20. Alkan C, Coe BP, Eichler EE: APPLICATIONS OF NEXT-GENERATION SEQUENCING Genome structural variation discovery and genotyping. Nature Reviews Genetics 2011, 12:363–375.

21. Ko BJ, Lee C, Kim J, Rhie A, Yoo DA, Howe K, Wood J, Cho S, Brown S, Formenti G, et al: Widespread false gene gains caused by duplication errors in genome assemblies. Genome Biol 2022, 23:205.

22. Nurk S, Koren S, Rhie A, Rautiainen M, Bzikadze AV, Mikheenko A, Vollger MR, Altemose N, Uralsky L, Gershman A, et al: The complete sequence of a human genome. Science 2022, 376:44–53.

23. Koren S, Rhie A, Walenz BP, Dilthey AT, Bickhart DM, Kingan SB, Hiendleder S, Williams JL, Smith TPL, Phillippy AM: **De novo assembly of haplotype-resolved genomes with trio binning**. Nature Biotechnology 2018, 36:1174-+.

24. Frantz LAF, Mullin VE, Pionnier-Capitan M, Lebrasseur O, Ollivier M, Perri A, Linderholm A, Mattiangeli V, Teasdale MD, Dimopoulos EA, et al: Genomic and archaeological evidence suggests a dual origin of domestic dogs. Science 2016, 352:1228–1231.

25. Lindblad-Toh K, Wade CM, Mikkelsen TS, Karlsson EK, Jaffe DB, Kamal M, Clamp M, Chang JL, Kulbokas EJ, 3rd, Zody MC, et al: **Genome sequence, comparative analysis and haplotype structure of the domestic dog**. Nature 2005, 438:803–819.

26. Hoeppner MP, Lundquist A, Pirun M, Meadows JRS, Zamani N, Johnson J, Sundström G, Cook A, FitzGerald MG, Swofford R, et al: An Improved Canine Genome and a Comprehensive Catalogue of Coding Genes and Non-Coding Transcripts. PLOS ONE 2014, 9:e91172.

27. Nicholas TJ, Cheng Z, Ventura M, Mealey K, Eichler EE, Akey JM: The genomic architecture of segmental duplications and associated copy number variants in dogs. Genome Research 2009, 19:491–499.

28. Chen WK, Swartz JD, Rush LJ, Alvarez CE: **Mapping DNA structural variation in dogs**. Genome Res 2009, 19:500–509.

29. Binversie EE, Baker LA, Engelman CD, Hao Z, Moran JJ, Piazza AM, Sample SJ, Muir P: Analysis of copy number variation in dogs implicates genomic structural variation in the development of anterior cruciate ligament rupture. PLoS One 2020, 15:e0244075.

30. Serres-Armero A, Davis BW, Povolotskaya IS, Morcillo-Suarez C, Plassais J, Juan D, Ostrander EA, Marques-Bonet T: **Copy number variation underlies complex phenotypes in domestic dog breeds and other canids**. Genome Res 2021, 31:762–774.

31. Pendleton AL, Shen F, Taravella AM, Emery S, Veeramah KR, Boyko AR, Kidd JM: Comparison of village dog and wolf genomes highlights the role of the neural crest in dog domestication. BMC Biology 2018, 16:64.

32. Serres-Armero A, Povolotskaya IS, Quilez J, Ramirez O, Santpere G, Kuderna LFK, Hernandez-Rodriguez J, Fernandez-Callejo M, Gomez-Sanchez D, Freedman AH, et al: Similar genomic proportions of copy number variation within gray wolves and modern dog breeds inferred from whole genome sequencing. BMC Genomics 2017, 18:977.

33. Wang C, Wallerman O, Arendt ML, Sundstrom E, Karlsson A, Nordin J, Makelainen S, Pielberg GR, Hanson J, Ohlsson A, et al: A novel canine reference genome resolves genomic architecture and uncovers transcript complexity. Commun Biol 2021, 4:185.

34. Sinding MS, Gopalakrishnan S, Raundrup K, Dalen L, Threlfall J, Darwin Tree of Life Barcoding c, Wellcome Sanger Institute Tree of Life p, Wellcome Sanger Institute Scientific Operations DNAPc, Tree of Life Core Informatics c, Darwin Tree of Life C, Gilbert T: **The genome sequence of the grey wolf, Canis lupus Linnaeus** 1758. Wellcome Open Res 2021, 6:310.

35. Player RA, Forsyth ER, Verratti KJ, Mohr DW, Scott AF, Bradburne CE: A novel canis lupus familiaris reference genome improves variant resolution for use in breed-specific GWAS. Life Science Alliance 2021, 4.

36. Jagannathan V, Hitte C, Kidd JM, Masterson P, Murphy TD, Emery S, Davis B, Buckley RM, Liu Y-H, Zhang X-Q, et al: Dog10K_Boxer_Tasha_1.0: A Long-Read Assembly of the Dog Reference Genome. Genes 2021, 12:847.

37. Halo JV, Pendleton AL, Shen F, Doucet AJ, Derrien T, Hitte C, Kirby LE, Myers B, Sliwerska E, Emery S, et al: Long-read assembly of a Great Dane genome highlights the contribution of GC-rich sequence and mobile elements to canine genomes. Proceedings of the National Academy of Sciences 2021, 118.

38. Field MA, Rosen BD, Dudchenko O, Chan EKF, Minoche AE, Edwards RJ, Barton K, Lyons RJ, Tuipulotu DE, Hayes VM, et al: Canfam_GSD: De novo chromosome-length genome assembly of the German Shepherd Dog (Canis lupus familiaris) using a combination of long reads, optical mapping, and Hi-C. Gigascience 2020, 9.

39. Field MA, Yadav S, Dudchenko O, Esvaran M, Rosen BD, Skvortsova K, Edwards RJ, Keilwagen J, Cochran BJ, Manandhar B, et al: The Australian dingo is an early offshoot of modern breed dogs. Sci Adv 2022, 8:eabm5944.

40. Edwards RJ, Field MA, Ferguson JM, Dudchenko O, Keilwagen J, Rosen BD, Johnson GS, Rice ES, Hillier LD, Hammond JM, et al: Chromosome-length genome assembly and structural variations of the primal Basenji dog (Canis lupus familiaris) genome. BMC Genomics 2021, 22:188.

41. Shen F, Kidd JM: Rapid, Paralog-Sensitive CNV Analysis of 2457 Human Genomes Using QuicK- mer2. Genes 2020, 11:141.

42. Wang G-D, Larson G, Kidd JM, vonHoldt BM, Ostrander EA, Zhang Y-P: **Dog10K: the International Consortium of Canine Genome Sequencing**. National Science Review 2019, 6:611–613.

43. Kent WJ, Sugnet CW, Furey TS, Roskin KM, Pringle TH, Zahler AM, Haussler D: **The human genome browser at UCSC**. Genome Res 2002, 12:996–1006.

44. Gene Ontology C, Aleksander SA, Balhoff J, Carbon S, Cherry JM, Drabkin HJ, Ebert D, Feuermann M, Gaudet P, Harris NL, et al: **The Gene Ontology knowledgebase in** 2023. Genetics 2023, 224.

45. Ashburner M, Ball CA, Blake JA, Botstein D, Butler H, Cherry JM, Davis AP, Dolinski K, Dwight SS, Eppig JT, et al: Gene ontology: tool for the unification of biology. The Gene Ontology Consortium. Nat Genet 2000, 25:25–29.

46. Thomas PD, Ebert D, Muruganujan A, Mushayahama T, Albou LP, Mi H: PANTHER: Making genome-scale phylogenetics accessible to all. Protein Sci 2022, 31:8–22.

47. Szak ST, Pickeral OK, Makalowski W, Boguski MS, Landsman D, Boeke JD: **Molecular archeology of L1 insertions in the human genome**. Genome Biol 2002, 3:research0052.

48. Feng Q, Moran JV, Kazazian HH, Jr., Boeke JD: Human L1 retrotransposon encodes a conserved endonuclease required for retrotransposition. Cell 1996, 87:905–916.

49. Flasch DA, Macia A, Sanchez L, Ljungman M, Heras SR, Garcia-Perez JL, Wilson TE, Moran JV: **Genome-wide de novo L1 Retrotransposition Connects Endonuclease Activity with Replication**. Cell 2019, 177:837–851 e828.

50. Jurka J: Sequence patterns indicate an enzymatic involvement in integration of mammalian retroposons. Proc Natl Acad Sci U S A 1997, 94:1872-1877.

51. Morrish TA, Gilbert N, Myers JS, Vincent BJ, Stamato TD, Taccioli GE, Batzer MA, Moran JV: **DNA repair mediated by endonuclease-independent LINE-1 retrotransposition**. Nat Genet 2002, 31:159–165.

52. Gao X, Li Y, Adetula AA, Wu Y, Chen H: **Analysis of new retrogenes provides insight into dog adaptive evolution**. Ecology and Evolution 2019, 9:11185–11197.

53. Liu YJ, Zheng D, Balasubramanian S, Carriero N, Khurana E, Robilotto R, Gerstein MB: Comprehensive analysis of the pseudogenes of glycolytic enzymes in vertebrates: the anomalously high number of GAPDH pseudogenes highlights a recent burst of retrotrans- positional activity. BMC Genomics 2009, 10:480.

54. Breen M: Canine cytogenetics--from band to basepair. Cytogenet Genome Res 2008, 120:50-60.

55. Marquez LM, Miller DJ, MacKenzie JB, Van Oppen MJ: Pseudogenes contribute to the extreme diversity of nuclear ribosomal DNA in the hard coral Acropora. Mol Biol Evol 2003, 20:1077–1086.

56. Ciomborowska-Basheer J, Staszak K, Kubiak MR, Makalowska I: Not So Dead Genes-Retrocopies as Regulators of Their Disease-Related Progenitors and Hosts. Cells 2021, 10.

57. Kukekova AV, Johnson JL, Xiang X, Feng S, Liu S, Rando HM, Kharlamova AV, Herbeck Y, Serdyukova NA, Xiong Z, et al: Red fox genome assembly identifies genomic regions associated with tame and aggressive behaviours. Nat Ecol Evol 2018, 2:1479–1491.

58. Wang GD, Shao XJ, Bai B, Wang J, Wang X, Cao X, Liu YH, Wang X, Yin TT, Zhang SJ, et al: Structural variation during dog domestication: insights from gray wolf and dhole genomes. Natl Sci Rev 2019, 6:110–122.

59. Koch E, Schweizer RM, Schweizer TM, Stahler DR, Smith DW, Wayne RK, Novembre J: **De novo mutation rate estimation in wolves of known pedigree**. Mol Biol Evol 2019, 36:2536–2547.

60. Linardopoulou EV, Williams EM, Fan Y, Friedman C, Young JM, Trask BJ: Human subtelomeres are hot spots of interchromosomal recombination and segmental duplication. Nature 2005, 437:94–100.

61. Sulak M, Fong L, Mika K, Chigurupati S, Yon L, Mongan NP, Emes RD, Lynch VJ: TP53 copy number expansion is associated with the evolution of increased body size and an enhanced DNA damage response in elephants. Elife 2016, 5.

62. Batcher K, Varney S, Raudsepp T, Jevit M, Dickinson P, Jagannathan V, Leeb T, Bannasch D: **Ancient segmentally duplicated LCORL retrocopies in equids**. PLoS One 2023, 18:e0286861.

63. Segurel L, Wyman MJ, Przeworski M: **Determinants of mutation rate variation in the human germline**. Annu Rev Genomics Hum Genet 2014, 15:47–70.

64. Lipson M, Loh PR, Sankararaman S, Patterson N, Berger B, Reich D: **Calibrating the Human Mutation Rate via Ancestral Recombination Density in Diploid Genomes**. PLoS Genet 2015, 11:e1005550.

65. Narasimhan VM, Rahbari R, Scally A, Wuster A, Mason D, Xue Y, Wright J, Trembath RC, Maher ER, van Heel DA, et al: Estimating the human mutation rate from autozygous segments reveals population differences in human mutational processes. Nat Commun 2017, 8:303.

66. Palamara PF, Francioli LC, Wilton PR, Genovese G, Gusev A, Finucane HK, Sankararaman S, Genome of the Netherlands C, Sunyaev SR, de Bakker PI, et al: **Leveraging Distant Relatedness to Quantify Human Mutation and Gene-Conversion Rates**. Am J Hum Genet 2015, 97:775–789.

67. Tian X, Browning BL, Browning SR: Estimating the Genome-wide Mutation Rate with Three- Way Identity by Descent. Am J Hum Genet 2019, 105:883–893.

68. Skoglund P, Ersmark E, Palkopoulou E, Dalen L: Ancient wolf genome reveals an early divergence of domestic dog ancestors and admixture into high-latitude breeds. Curr Biol 2015, 25:1515–1519.

69. Bergstrom A, Frantz L, Schmidt R, Ersmark E, Lebrasseur O, Girdland-Flink L, Lin AT, Stora J, Sjogren KG, Anthony D, et al: Origins and genetic legacy of prehistoric dogs. Science 2020, 370:557–564.

70. Botigue LR, Song S, Scheu A, Gopalan S, Pendleton AL, Oetjens M, Taravella AM, Seregely T, Zeeb-Lanz A, Arbogast RM, et al: Ancient European dog genomes reveal continuity since the Early Neolithic. Nat Commun 2017, 8:16082.

71. Fan Z, Silva P, Gronau I, Wang S, Armero AS, Schweizer RM, Ramirez O, Pollinger J, Galaverni M, Ortega Del-Vecchyo D, et al: Worldwide patterns of genomic variation and admixture in gray wolves. Genome Res 2016, 26:163–173.

72. Sinding MS, Gopalakrishan S, Vieira FG, Samaniego Castruita JA, Raundrup K, Heide Jorgensen MP, Meldgaard M, Petersen B, Sicheritz-Ponten T, Mikkelsen JB, et al: Population genomics of grey wolves and wolf-like canids in North America. PLoS Genet 2018, 14:e1007745.

73. Ewing AD, Ballinger TJ, Earl D, Broad Institute Genome S, Analysis P, Platform, Harris CC, Ding L, Wilson RK, Haussler D: **Retrotransposition of gene transcripts leads to structural variation in mammalian genomes**. Genome Biol 2013, 14:R22.

74. Nachman MW, Crowell SL: Estimate of the mutation rate per nucleotide in humans. Genetics 2000, 156:297–304.

75. Jonsson H, Sulem P, Kehr B, Kristmundsdottir S, Zink F, Hjartarson E, Hardarson MT, Hjorleifsson KE, Eggertsson HP, Gudjonsson SA, et al: Parental influence on human germline de novo mutations in 1,548 trios from Iceland. Nature 2017, 549:519–522.

76. Feng X, Li H: Higher Rates of Processed Pseudogene Acquisition in Humans and Three Great Apes Revealed by Long-Read Assemblies. Mol Biol Evol 2021, 38:2958–2966.

77. Batcher K, Dickinson P, Maciejczyk K, Brzeski K, Rasouliha SH, Letko A, Drogemuller C, Leeb T, Bannasch D: **Multiple FGF4 Retrocopies Recently Derived within Canids**. Genes (Basel) 2020, 11.

78. Wang W, Kirkness EF: Short interspersed elements (SINEs) are a major source of canine genomic diversity. Genome Res 2005, 15:1798–1808.

79. Beck CR, Garcia-Perez JL, Badge RM, Moran JV: **LINE-1 elements in structural variation and disease**. Annu Rev Genomics Hum Genet 2011, 12:187–215.

80. Wei W, Gilbert N, Ooi SL, Lawler JF, Ostertag EM, Kazazian HH, Boeke JD, Moran JV: **Human L1 retrotransposition: cis preference versus trans complementation**. Mol Cell Biol 2001, 21:1429–1439.

81. Dewannieux M, Esnault C, Heidmann T: **LINE-mediated retrotransposition of marked Alu sequences**. Nat Genet 2003, 35:41–48.

82. Verscheure S, Backeljau T, Desmyter S: In silico discovery of a nearly complete mitochondrial genome Numt in the dog (Canis lupus familiaris) nuclear genome. Genetica 2015, 143:453–458.

83. Quinlan AR: BEDTools: The Swiss-Army Tool for Genome Feature Analysis. Curr Protoc Bioinformatics 2014, 47:11 12 11-34.

84. Benson G: Tandem repeats finder: a program to analyze DNA sequences. Nucleic Acids Res 1999, 27:573-580.

85. Morgulis A, Gertz EM, Schaffer AA, Agarwala R: **WindowMasker: window-based masker for sequenced genomes**. Bioinformatics 2006, 22:134–141.

86. Hach F, Hormozdiari F, Alkan C, Hormozdiari F, Birol I, Eichler EE, Sahinalp SC: **mrsFAST: a cache- oblivious algorithm for short-read mapping**. Nat Methods 2010, 7:576–577.

87. Li H: Minimap2: pairwise alignment for nucleotide sequences. Bioinformatics 2018, 34:3094- 3100.

88. Shumate A, Salzberg SL: **Liftoff: accurate mapping of gene annotations**. Bioinformatics 2021, 37:1639–1643.

89. Danecek P, Bonfield JK, Liddle J, Marshall J, Ohan V, Pollard MO, Whitwham A, Keane T, McCarthy SA, Davies RM, Li H: **Twelve years of SAMtools and BCFtools**. Gigascience 2021, 10.

90. Kent WJ: BLAT--the BLAST-like alignment tool. Genome Res 2002, 12:656-664.

91. Li H: New strategies to improve minimap2 alignment accuracy. Bioinformatics 2021, 37:4572- 4574.

92. Rice P, Longden I, Bleasby A: EMBOSS: the European Molecular Biology Open Software Suite. Trends Genet 2000, 16:276–277.

93. Smith TF, Waterman MS: Identification of common molecular subsequences. J Mol Biol 1981, 147:195–197.

94. Hunter JD: Matplotlib: A 2D Graphics Environment. Computing in Science & Engineering 2007, 9:90–95.

95. Tareen A, Kinney JB: Logomaker: beautiful sequence logos in Python. Bioinformatics 2020, 36:2272–2274.

96. Lex A, Gehlenborg N, Strobelt H, Vuillemot R, Pfister H: UpSet: Visualization of Intersecting Sets. IEEE Trans Vis Comput Graph 2014, 20:1983–1992.

97. Freedman AH: Genome Sequencing Highlights the Dynamic Early History of Dogs (vol 10, e1004016, 2014). Plos Genetics 2014, **10**.

98. Larson G, Bradley DG: How Much Is That in Dog Years? The Advent of Canine Population Genomics. Plos Genetics 2014, 10.

